# Bayesian Estimation of Species Divergence Times Using Correlated Quantitative Characters

**DOI:** 10.1101/441105

**Authors:** Sandra Álvarez-Carretero, Anjali Goswami, Ziheng Yang, Mario dos Reis

## Abstract

Discrete morphological data have been widely used to study species evolution, but the use of quantitative (or continuous) morphological characters is less common. Here, we implement a Bayesian method to estimate species divergence times using quantitative characters. Quantitative character evolution is modelled using Brownian diffusion with character correlation and character variation within populations. Through simulations, we demonstrate that ignoring the population variation (or population “noise”) and the correlation among characters leads to biased estimates of divergence times and rate, especially if the correlation and population noise are high. We apply our new method to the analysis of quantitative characters (cranium landmarks) and molecular data from carnivoran mammals. Our results show that time estimates are affected by whether the correlations and population noise are accounted for or ignored in the analysis. The estimates are also affected by the type of data analysed, with analyses of morphological characters only, molecular data only, or a combination of both; showing noticeable differences among the time estimates. Rate variation of morphological characters among the carnivoran species appears to be very high, with Bayesian model selection indicating that the independent-rates model fits the morphological data better than the autocorrelated-rates model. We suggest that using morphological continuous characters, together with molecular data, can bring a new perspective to the study of species evolution. Our new model is implemented in the MCMCtree computer program for Bayesian inference of divergence times. [Bayesian inference, continuous morphological characters, geometric morphometrics, Procrustes alignment, molecular clock, divergence times, phylogeny]

---

Molecular sequences are informative about the relative ages of nodes on a phylogeny, but not about the geological times of divergence or the absolute molecular evolutionary rate. The Bayesian method offers a way to use fossil information to construct a prior on divergence times, which can then be integrated with the molecular data to produce posterior estimates of absolute divergence times (e.g., Thorne et al., 1998; Drummond et al., 2006; Rannala and Yang, 2007). However, modelling clade ages with statistical distributions based on the fossil evidence is challenging. Fossils may provide estimates of minimum clade ages, but maximum clade ages are typically based on the absence of fossil evidence, and are thus hard to justify (Benton and Donoghue, 2007).

The problem is illustrated in Figure 1. Imagine we wish to estimate the age of the last common ancestor of species A and B, *t*_*AB*_. The oldest fossil in the A-B ingroup is F, which has known age *t*_*F*_. If we measure time towards the past (so that present time is zero), we can immediately see that *t_AB_ > t_F_*, so that the age of the fossil, *t*_*F*_, imposes a minimum constraint on *t*_*AB*_. However, we do not know how close F is to the common ancestor, so *t*_*F*_ is a poor indicator of the true age *t*_*AB*_. Current practice is to construct a prior density on *t*_*AB*_, *f* (*t*_*AB*_), truncated at *t*_*F*_ on the left, and with a long tail extending to the right (back in time) to allow for the uncertainty in the time gap between *t*_*F*_ and *t*_*AB*_ (Fig. 1). The form of the prior density and the length of the tail are somewhat subjective as they are based on absence of older fossils in the A-B clade (e.g., Tavaré et al., 2002; Drummond et al., 2006; Yang and Rannala, 2006; Benton and Donoghue, 2007).

**Figure 1:**
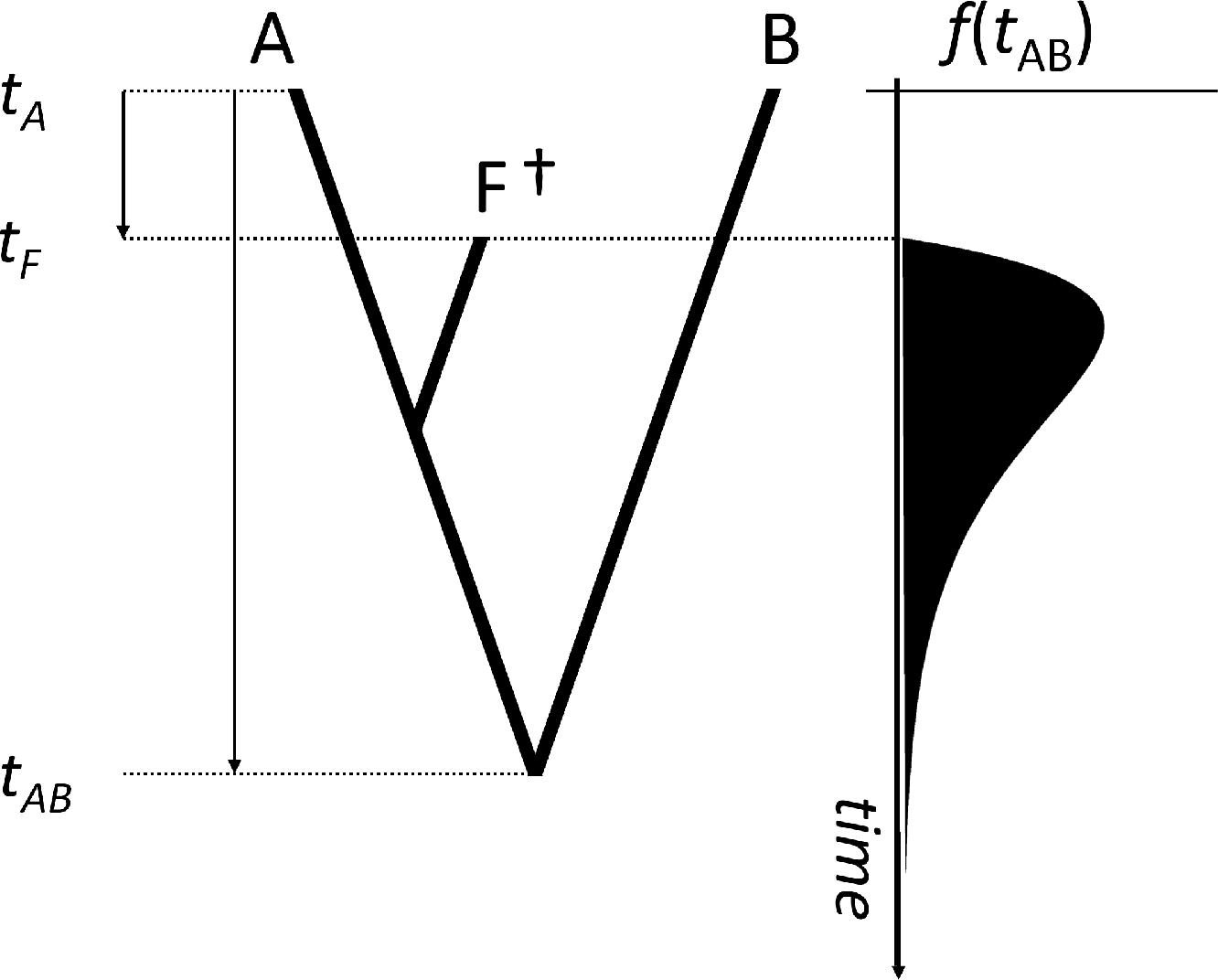
A phylogeny of two extant species (A and B) and one extinct species (F*†*). The age of the extinct fossil, *t*_*F*_, provides a minimum age bound on the divergence of A and B, *t*_*AB*_. The fossil age can be used as a lower limit on a prior probability distribution, *f* (*t*_*AB*_), in a Bayesian analysis. Deciding on the shape of the distribution and on how far its tail should stretch back in time is somewhat subjective (Donoghue and Benton, 2007). Here we show an example of a misspecified prior for *t*_*AB*_, with the probability mass close to the age of the fossil, but too far from the true age of the node.

An alternative approach would be to model morphological character evolution, so that we can use morphological data to estimate the morphological distance among extant and fossil species in a phylogeny. Since fossil ages are known, fossils can then be used as “dated-tips” in the Bayesian analysis. Divergence time estimation can then proceed using a morphological alignment of extant and fossil species, or on a combined data set of molecular data for extant species and morphological data for extant and fossil species. This approach, also known as total-evidence dating (TED), has been pioneered by Pyron (2011) and Ronquist et al. (2012) (see also Nylander et al., 2004; Lee et al., 2009; and Magallón, 2010) using discrete morphological characters under the Mk model of morphological evolution (Lewis, 2001). It has been used to date phylogenies for several groups (e.g., Nylander et al., 2004; Pyron, 2011; Ronquist et al., 2012; Schrago et al., 2013; Slater, 2013; Wood et al., 2013; Arcila et al., 2015; Grimm et al., 2015; Reeder et al., 2015; Winterton and Ware, 2015; Larson-Johnson, 2016; Ronquist et al., 2016; Gavryushkina et al., 2017), sometimes producing very old time estimates compared with node-calibration methods, and it is noted to be sensitive to the branching process used to specify the prior on times (O’Reilly et al., 2015; dos Reis et al., 2016). The TED approach has been improved by extensions of the fossilised birth-death process to construct more realistic priors on times (Heath et al., 2014; Gavryushkina et al., 2014; Zhang et al., 2016).

Analysis of discrete morphological data under the Mk model has a few limitations. First, the model assumes that rates of change among character states are equal (Lewis, 2001), an assumption that appears unrealistic for the analysis of real data. Although the equal-rates assumption can be relaxed (Pagel, 1994; Wright et al., 2016), this model appears to be rarely used, perhaps because it is computationally expensive (Wright et al., 2016). Second, systematists usually score discrete morphological characters only if the characters are variable or if they are parsimony-informative. In this case, a correction is necessary to account for the ascertainment bias in character scoring (Lewis, 2001; Leaché et al., 2015). Correcting for the removal of constant characters is straightforward, but a much more computationally expensive correction is necessary to account for the removal of parsimony-uninformative characters, and it appears that this correction is not properly accommodated in current dating software (dos Reis et al., 2016). Finally, it seems difficult to accommodate correlations among characters in the Mk model. For a morphological alignment with *p* characters and with each character having *k* states, a *k*^*p*^ substitution matrix is constructed to accommodate correlated character evolution (Pagel, 1994). Such matrices become explosively large for even a moderate number of characters and are computationally intractable (Felsenstein, 2005). Thus, correlation among characters is ignored in Bayesian inference under the Mk model. The threshold model, an alternative to the Mk model for the analysis of ordered categorical data that may easily accommodate correlations among characters, has been championed by Felsenstein (2005; 2012). However, this model does not appear to be currently available for Bayesian inference of topology or divergence times of phylogenies.

Quantitative (or continuous) morphological characters offer an interesting alternative to the analysis of discrete characters (Felsenstein, 1988; Slater et al., 2012; Parins-Fukuchi, 2018b,a). Evolution of quantitative characters on a phylogeny can be modelled using diffusion processes such as the Brownian or Ornstein-Uhlenbeck processes (Felsenstein 1973; 1988). An appealing property of these processes is that the resulting likelihood of the characters on the phylogeny is a multivariate normal distribution which can be extended to accommodate correlations among characters and can be easily calculated. Furthermore, because quantitative characters are always variable, an ascertainment bias correction is not necessary. Also, non-homogeneity among characters can be easily accommodated in the normal likelihood: each character may have its own diffusion rate and its own ancestral mean, and thus expensive integration over a distribution of stationary frequencies (as done for the relaxed version of the Mk model, see Wright et al., 2016) is not necessary.

Here we explore the use of quantitative characters for Bayesian inference of species divergence times under the Brownian diffusion model of Felsenstein (1973). We use computer simulations to study the performance of the model in obtaining divergence time estimates: we focus on the effect of the sample size (the number of characters analysed) and the fossil age (using young or old fossils in the phylogeny), the strength of the correlation among the characters, and the level of “population noise” on the performance of the method. In the Brownian diffusion model, the means of the characters in populations evolve according to Brownian diffusion, but quantitative measurements on a sample of individuals for a given population of species is expected to show variation around the population mean. This population noise must be explicitly accommodated in the model (Felsenstein, 1973). Finally, we study the performance of the method on the analysis of a real data set: a set of cranium landmarks on a carnivoran phylogeny.

## Theory

We assume that the species phylogeny (the tree topology) is known. The posterior distribution of times and rates is

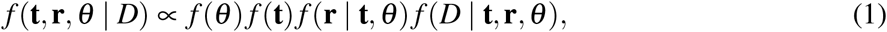

where *f* (*θ*) is the prior on model parameters, *f* (**t**) is the prior on times, *f* (**r**| **t**, *θ*) is the prior on rates, and *f* (*D* | **t**, **r**, *θ*) is the likelihood of the data *D*. In this paper, the data *D* may be a molecular sequence alignment *S*, a morphological alignment *M*, or a combination of both. Evolutionary rates may then include molecular rates **r**_*S*_ and/or morphological rates **r**_*M*_. For combined data, and assumming independent evolution of molecular and morphological characters, the posterior is

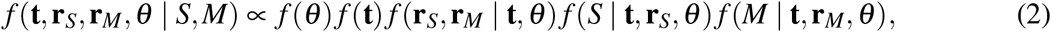

where *f* (*S* | **t**, **r**_*S*_, *θ*) is the likelihood of the molecular sequence alignment (e.g., calculated under the HKY+Γ substitution model) and *f* (*M* | **t**, **r**_*M*_, *θ*) is the likelihood of the morphological alignment, calculated under the Brownian diffusion model of quantitative character evolution (Felsenstein, 1973).

### Likelihood Calculation of Quantitative Characters

Calculation of the likelihood is described by Felsenstein (1973; 1981; see also Freckleton, 2012). Let **M** = {*m*_*i j*_} be a matrix of *p* continuous morphological characters measured on *s* species, where *m_i j_* is the *j*-th morphological measurement in species *i*, with *i* = 1, …, *s* and *j* = 1, …, *p*. Let **m**_*i*_ be the vector of *p* measurements in species *i* (the *i*-th row of **M**). Let **R** be the *p*× *p* correlation matrix among the characters. Write **m**_*s*+1_ for the vector of *p* (unobserved) ancestral character states at the root of the phylogeny. Character *j* evolves from its ancestral state *m*_*s*+1, *j*_ to a terminal state *m*_*i, j*_ along the branches of the tree by Brownian motion with diffusion rate *r* = *σ*^2^ (where *σ* is the diffusion coefficient, Felsenstein, 1973). Then, *m*_*i, j*_ is normally distributed with mean *m*_*s*+1, *j*_ and variance *v* = *rt*, where *t* is the elapsed time between the root and the tip species. If we assume that the rates (and thus the variances) are the same across characters (an assumption that can be relaxed), then **m**_*i*_ has a multivariate normal distribution with mean **m**_*s*+1_ and covariance matrix *v***R**. The diffusion rates may vary among lineages (branches) in a phylogeny (Felsenstein, 1981). If *r_k_* is the rate in branch *k*, and *t_k_* is the elapsed time along the branch, then *v_k_* = *r*_*k*_*t*_*k*_ is the expected amount of morphological variance accumulated in the lineage. Thus *v*_*k*_ is the morphological branch length. Felsenstein (1973) showed that the likelihood of **M** on a phylogeny of two or more species can be calculated so that it only depends on the branch lengths, **v** = (*v*_*k*_), and the correlation matrix, **R**, but not on the ancestral characters at the root, **m**_*s*+1_. This simplifies the calculations as **m**_*s*+1_ does not need to be estimated.

Now consider a bifurcating, rooted phylogeny of *s* species. The external nodes (the tips) are labelled 1, …, *s*; the internal nodes are labelled *s* + 1, …, 2*s* − 1; and *s* + 1 is the root node. The length of the branch subtending node *k* is *v*_*k*_. If *k* is an internal node, let *k*_1_ and *k*_2_ be its two daughter nodes. Let

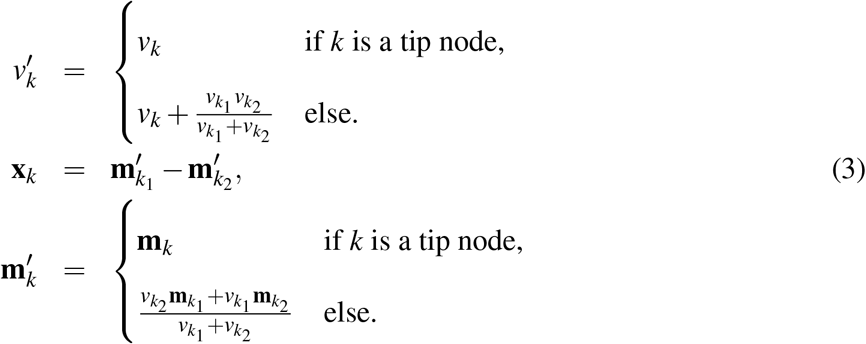

The likelihood of **M** on the phylogeny is the product of *s* − 1 multivariate normal densities, each corresponding to one of the *s* − 1 internal nodes. It is given by

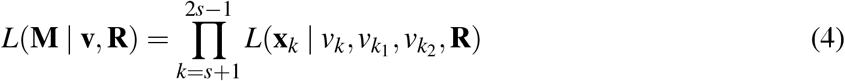

where

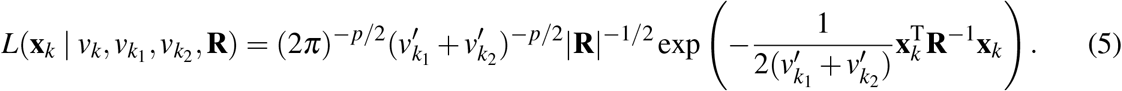

Eq. (4) can be calculated efficiently in a computer program using the postorder tree traversal algorithm. When an internal node *k* is visited by the algorithm, we calculate 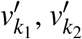, **x**_*k*_ and 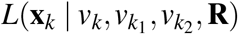 after its daughter nodes have been visited. The **m**′_*k*_ are maximum likelihood estimates of the ancestral character states at node *k* conditioned on the values of 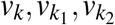, and **R**. They are obtained for free during MCMC computation, and they may be collected and used as ancestral reconstructions.

### Correlation Among Characters and Matrix Shrinkage

It is useful to find a matrix **A** such that **R**^−1^ = **A**^T^**A**. Then, the exponential in the likelihood of Eq. (5) can be written as

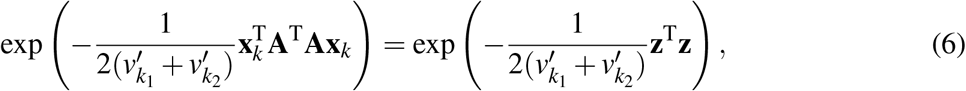

where **z** = **Ax**_*k*_ is a vector. In other words, we can obtain a transformation of the original data **Z** = **MA**^T^, so that the transformed characters in **Z** are independent. This simplifies the calculation of the likelihood because **R** only needs to be inverted/decomposed once. Choices for **A** include the Cholesky decomposition, **R** = **LL**^T^, then **A** = **L**^−1^, or the Eigen decomposition **A**^T^ = **VD**, where **V** is the matrix of eigenvectors of **R**^−1^, and 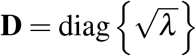 is a diagonal matrix of the square root of the eigenvalues (see p. 98 in Ripley, 1987).

The correlation matrix **R** can be estimated during Bayesian inference. However, this would make computation prohibitively expensive as we would need to estimate (*p*^2^ − *p*)/2 correlations, which is a large number for even a moderate *p*. Thus, here we assume that **R** is given. For example, if we assume **R** is constant throughout the phylogeny, then we can estimate **R** from a sample of individuals from a given species. The individuals may be assumed to be independent samples from the population, and **R** could then be estimated using the traditional unbiased estimate of the covariance. However, a common problem occurs when the number of characters, *p*, is larger than the number of individuals sampled, *s*. In this case, the unbiased estimate of **R**, **R̂**, tends to become singular (i.e., its determinant is zero) and cannot be inverted (e.g., Schäfer and Strimmer, 2005; Goolsby, 2016), in which case the likelihood of Eq. (5) cannot be calculated. Here we overcome this problem by using the linear shrinkage estimate of the correlation matrix (Schäfer and Strimmer, 2005):

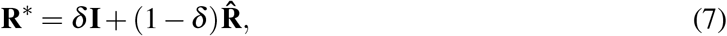

where **I** is the identity matrix, and *δ* (0 ≤ *δ* ≤ 1) is the shrinkage parameter, which controls the level of shrinkage. If *δ* = 0, the shrinkage estimate, **R**\*, is the same as **R̂**, while if *δ* = 1, **R**\* is the identity matrix.

Note that **R**\* can always be inverted as long as *δ* ≠ 0, thus allowing calculation of the likelihood of Eq. (5). The value of *δ* can be chosen by the user or estimated automatically. Schäfer and Strimmer (2005) give an approximate method for automatically estimating *δ* from the data. Their procedure is implemented in their `corpcor R` package (see their paper for details of the algorithm). Clavel et al. (2018) discuss further approaches to regularise the estimate of **R**.

### Within Population Character Variance

Quantitative characters are expected to vary among individuals within a species (Felsenstein, 1973; Ives et al., 2007). Divergence times may be biased if this population level variation (or “population noise”) is large and not accounted for in the Bayesian inference, because the amount of morphological evolution in the phylogeny would be overestimated (Landis and Schraiber, 2017). Write *c*_*j*_ for the within population variance of character *j*. Then *m*_*i, j*_ is normally distributed with mean *m*_*s*+1, *j*_ and variance *c*_*j*_ + *v*. In this case, *m*_*s*+1, *j*_ is then the population mean of the character in the ancestral population (Felsenstein, 1973). If all characters have the same variance *c*, then we can accommodate the population noise in the analysis by setting

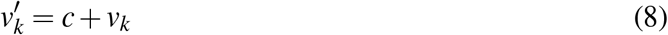

when *k* is a tip node (Eqs. 3 and 5).

In real data, different characters may have different variances. In this case, we may obtain estimates of the variances of the characters, **ĉ** = (*ĉ*_*j*_), from a population sample at the same time as we estimate **R**. We can then divide the columns of **M** by the estimated standard deviations to obtain the scaled matrix 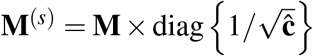. The new scaled matrix has thus been standardised so that all characters have the same variance and so that the population noise has unit variance. Inference then proceeds on **M**^(*s*)^ by setting *c* = 1 in Eq. (8). Note that to correct for the correlations among characters, the transformed data matrix used during Bayesian inference is then **Z**(*s*) = **M**(*s*)**A**^T^.

### Within-lineage and Among-lineage Covariances

We note that **R** here is the within-lineage correlation among the characters, and thus *v*_*k*_**R** is the within-lineage covariance for the *k*-th branch. For example, if selective pressure acts to elongate a limb in a species, one would expect the length of the corresponding limb bones to increase. In other words, the bone lengths would co-vary (or co-drift in Brownian parlance) and this would be represented by a positive correlation in the entry of **R** for the given characters. If the within-lineage variances are different among characters, then the exponent of Eq. (5) must be written as

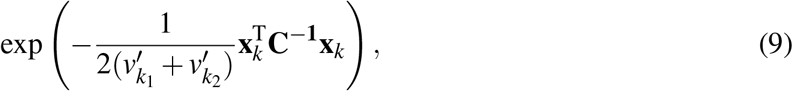

where **C** is then the within-lineage character covariance matrix (this is the same **C** as in Freckleton, 2012). However, if we can normalise the characters to have equal variances by using estimates of the within-population variances (as we do here and as shown in Felsenstein, 1973), then it is not necessary to work with the more complex Eq. (9).

The shared ancestry among the species in a phylogeny means that there is also character covariation among lineages. The among-lineage covariance matrix is *r***T** when *r* is constant (e.g., when we have a strict morphological clock), and where **T** is an *s* × *s* matrix whose elements are the shared ancestry time-paths for each pair of species (Felsenstein, 1973). For a Brownian model with unequal diffusion rates among branches (Felsenstein, 1981), we must multiply the shared time-paths in **T** by the branch-specific diffusion rates, *r*_*k*_, resulting in the *s* × *s* among-lineage covariance matrix **V**(see Felsenstein, 1981;Freckleton, 2012). Matrix **V** only appears explicitly when we write down the joint likelihood for the characters for all species (e.g., Eqs. 1 and 8 in Freckleton, 2012). Eq. (5) here is the node likelihood after the pruning algorithm has been applied, and thus matrix **V** is not apparent. However, note the *v*′_*k*_ terms are functions of the entries in **V**. See Felsenstein (1973; 1981) and Freckleton (2012) for full details.

## Software

Bayesian inference of divergence times with continuous characters under the model of Eq. (4) is implemented in the computer program MCMCtree v4.9i, part of the PAML package (Yang, 2007). We have extended the `mcmc3r R` package (dos Reis et al., 2018; https://github.com/dosreislab/mcmc3r) to help the user in preparing morphological alignments for analysis with MCMCtree, and in simulating continuous morphological data using different functions from the ape package (Paradis et al., 2004).

## Simulation Analysis

We conduct a simulation study to examine the impact of (i) the number of characters analysed, (ii) the fossil ages, (iii) the population noise, and (iv) the correlation among characters on the estimation of divergence times using morphological data. In particular, we expect that time estimates will deteriorate (i.e., will have large variances or be biased) when analysing small data numbers of characters, when the fossils are too young, when the population noise is high, and when the correlation among characters is strong. To reduce the computational cost, our simulations are carried out using a small number of species under a constant morphological evolutionary rate.

### Simulation overview

The phylogeny in Figure 2, with *s* = 8 species (5 extant and 3 fossils), is used to simulate the quantitative morphological data sets. The morphological evolutionary rate is *r* = 1 and constant along all the branches of the phylogeny. The simulated data matrices, **M**, are generated under the Brownian diffusion model using our `mcmc3r R` package. For simulations with population noise and/or correlations, a population sample of individuals is simulated, which is then used to estimate the vector of character variances, **ĉ**, and the shrinkage estimate of the correlation, **R**\*.

**Figure 2:**
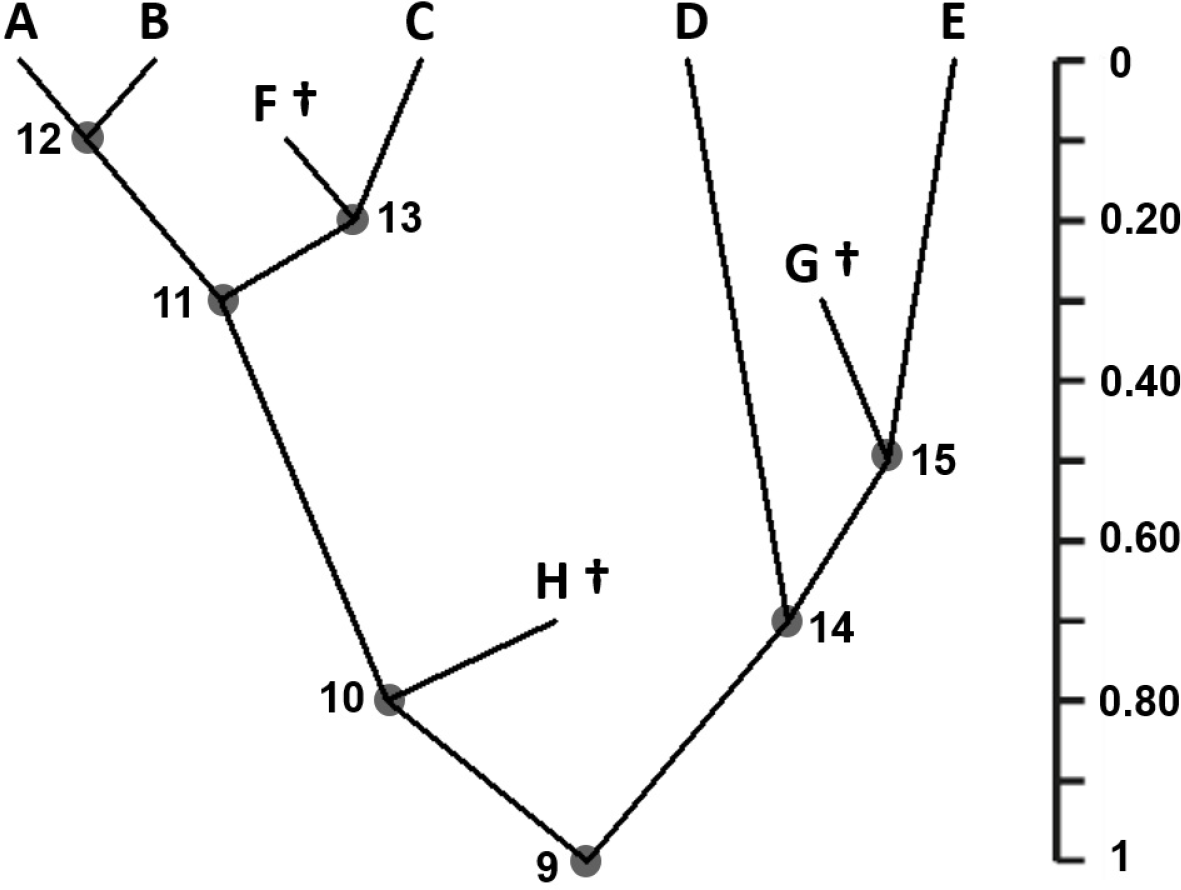
A phylogeny of 8 species used to simulate morphological data. The time unit is 100 myr and the divergence times are: *t*_9_ = 1.0 (root), *t*_10_ = 0.8, *t*_11_ = 0.3, *t*_12_ = 0.1, *t*_13_ = 0.2, *t*_14_ = 0.7, and *t*_15_ = 0.5; meaning 100 Ma, 80 Ma, 30 Ma, and so on. The ages of the fossils are *t*_*F*_ = 0.1, *t*_*G*_ = 0.3 and *t*_*H*_ = 0.7. Fossil species are indicated with a dagger (*†*).

Replicates under each simulation setup (see below) are analysed with MCMCtree to estimate the divergence times (*t*_9_ to *t*_15_) and the morphological rate (*r*) by MCMC sampling. We use a diffuse gamma prior on the rate, *r* ~ Γ(2, 2), with mean 1 and variance 0.5. The prior on the age of the root is assigned a uniform density with soft bounds between 0.8 and 1.2 (corresponding to a calibration of 80 to 120 Ma given our 100 myr time unit). The birth-death-sequential-sampling (BDSS) process (Stadler and Yang, 2013), is used to generate the prior density for the ages of the internal nodes. The BDSS parameters are set as: *λ*_BDSS_ = 1 (the birth-rate), *μ*_BDSS_ = 1 (the death-rate), *ρ*_BDSS_ = 0 (the sampling fraction for extant species), and *ψ*_BDSS_ = 0.001 (the rate of fossil sampling). We chose these parameter values to generate a uniform density on the ages (Fig. S1). We summarise the results by calculating the mean times across the replicates, the mean 95% credibility intervals (CIs), the mean CI-width *w* (and relative CI-width *w*_*r*_ = *w*/*t*_*i*_), the coverage (the number of times the true node age falls within the 95% CI), the mean bias, and the mean squared error (MSE). Let *t̃*_*i, j*_ be the mean posterior age of node *i* for replicate *j*. The mean bias is 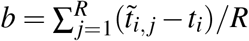 and the MSE is 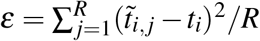, where *R* = 1, 000 is the number of replicates per simulation setup and *t*_*i*_ is the true node age. We also calculate the relative bias *b*_*r*_ = *b*/*t*_*i*_ and the relative error *∊*_*r*_ = *∊*/*t*_*i*_. Note the bias is a measure of accuracy of the estimate, while the MSE is a measure of both precision and accuracy. The simulation workflow is summarised in Figure S2.

i. *Effect of the number of characters*. – We simulate data sets with *p* = 100, 1, 000 and 10, 000 characters, assuming independence among characters and no population noise (*c* = 0).
ii. *Effect of fossil age*. – We vary the age of the fossil *H*, with *t*_*H*_ = 0.7, 0.5, 0.3, and 0.1. The ages of the other fossils remain unchanged. Characters are assumed to evolve independently and with *c* = 0. The data are then simulated using the phylogeny with the new fossil age with *p* = 100, 1, 000 and 10, 000 characters, respectively, giving 3 *×* 3 = 9 simulation setups.
iii. *Effect of population noise*. – We simulate data sets with *c* = 0.25 (low population noise) and *c* = 0.5 (high population noise) for *p* = 100, 1, 000 and 10, 000. Characters are assumed to evolve independently. In order to simulate the population noise, we sample *s× p* random numbers from a normal distribution with mean 0 and variance *c*, to obtain the *s× p* noise matrix **N**. The resulting noise is added to the simulated morphological matrix, **M**, to generate the noisy matrix **M**^(*n*)^ = **M**+ **N**. We also simulate a population sample of *n* = 20 individuals to obtain a *n× p* population matrix, **P**, by sampling from the normal distribution with mean 0 and variance *c*. Before performing Bayesian inference, we obtain estimates of the population noise for each character, **ĉ** = (*ĉ*_*j*_), using the simulated population sample **P**, and obtain the scaled matrix 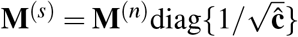. As we are scaling **M**^(*n*)^ using an estimate of the population variances, **ĉ**, we expect to observe little discrepancy between the true parameters (rate and divergence times) and their corresponding estimates. Therefore, we also scale the noisy matrix by **c** = (*c*_*j*_), the vector of true variances. Thus 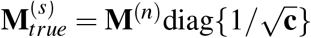, which is used as a control test. Bayesian inference then proceeds either on **M**^(*s*)^ or, 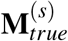 with the likelihood corrected by setting *c* = 1 (Eq. 8). The data are also analysed by setting *c* = 0 (Eq. 8) to assess the impact of ignoring the population noise on the time estimates. Note that the gamma prior on the morphological rate may be changed to account for scaling of the data sets. When *c* = 0.25, the morphological rate for the scaled data is *r/*0.25 = 1/0.25 = 4. Thus, the new gamma prior for the rate is *r* ~ Γ(2, 0.5). Similarly, when *c* = 0.5, the morphological rate for the scaled data is *r/*0.5 = 1/0.5 = 2, thus the rate prior is set to *r* ~ Γ(2, 1). We expect the posterior means of times and rates to be very biased when the population noise is ignored, to have some bias when using **M**^(*s*)^, and to have little or no bias when using _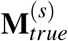_.
iv. *Effect of correlation among characters*. – We simulate data sets using the constant correlation model, that is, with all the within-lineage correlations in **R** equal to *ρ*. We use the correlations *ρ* = 0.5 and 0.9, and *p* = 100, 1, 000 and 10, 000. To simulate correlated data, a matrix **M** is first simulated assuming independent character evolution. Note that **M** is simulated on the tree, thus it already contains the among-lineage covariance induced by the shared ancestry. Then, we add the within-lineage correlation to **M** by computing **M**^(R)^ = **ML**^T^, where **L** is the lower triangular Cholesky decomposition of **R**. Then, we simulate the *s× p* noise matrix, **N**, sampled from a normal distribution with mean 0 and variance *c* = 0.25, to which within-lineage correlation is added as **N**^(R)^ = **NL**^T^. The noise is then added to **M**^(R)^ to obtain the noisy matrix, **M**^(*n*)^ = **M**^(R)^ + **N**^(R)^.

We also simulate a within-population sample of *n* = 20 individuals to obtain a *n× p* population matrix, **P**, by sampling from a normal distribution with mean 0 and variance *c* = 0.25, to which correlation is added as **P**^(R)^ = **PL**^T^. We use **P**^(R)^ to estimate **ĉ** = (*ĉ*_*j*_), the vector of estimated variances used to obtain **M**^(*s*)^. The vector of true variances, **c** = (*c*_*j*_), is used to obtain 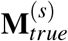. The shrinkage correlation matrix, **R**\*, is also estimated using **P**^(R)^. However, note that the shrinkage value, *δ*, has a strong impact on **R**\*. Therefore, we test two approaches to generate **R**\*: (i) we use the automatic approach of Schäfer and Strimmer (2005) to find the optimum value of *δ*, and (ii) we fix *δ* = 0.01, to obtain **R**\* close to the unbiased e stimate **R̂**. The Cholesky decomposition of **R**\* is then used to obtain the transformed data matrix **Z**^(*s*)^ = **M**^(*s*)^**A**^T^. **M**^(*s*)^ is also analysed directly to assess the effect of ignoring the character correlation. As a control data set, we also obtain **A** from the true correlation matrix, **R**, and use it to calculate 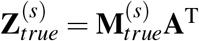. The estimates obtained using 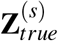 are expected to be very close to the true rate and divergence times. On the other hand, we expect estimates using **Z**^(*s*)^ to have some bias, and estimates using the uncorrected matrix, **M**^(*s*)^ (which ignores the correlation), to be very biased.

## Analysis of the Carnivora Data Set

### Morphological Data

We analyse the 29 cranium landmarks from 10 extant and 9 extinct carnivoran species (Fig. 3 and Table 1). The landmark data is complete (that is, there are no missing landmarks in any specimens). The landmarks are three dimensional, resulting in *p* = 3 *×* 29 = 87 characters. A population sample of 21 common foxes (*Vulpes vulpes*) is used to estimate the correlations and the population noise for the 29 landmarks. The correlation matrix estimated using the foxes is then used to transform the whole dataset (Eq. 6). This assumes the within-lineage correlations are the same (or at least similar) among the carnivorans analysed.

**Table 1:**
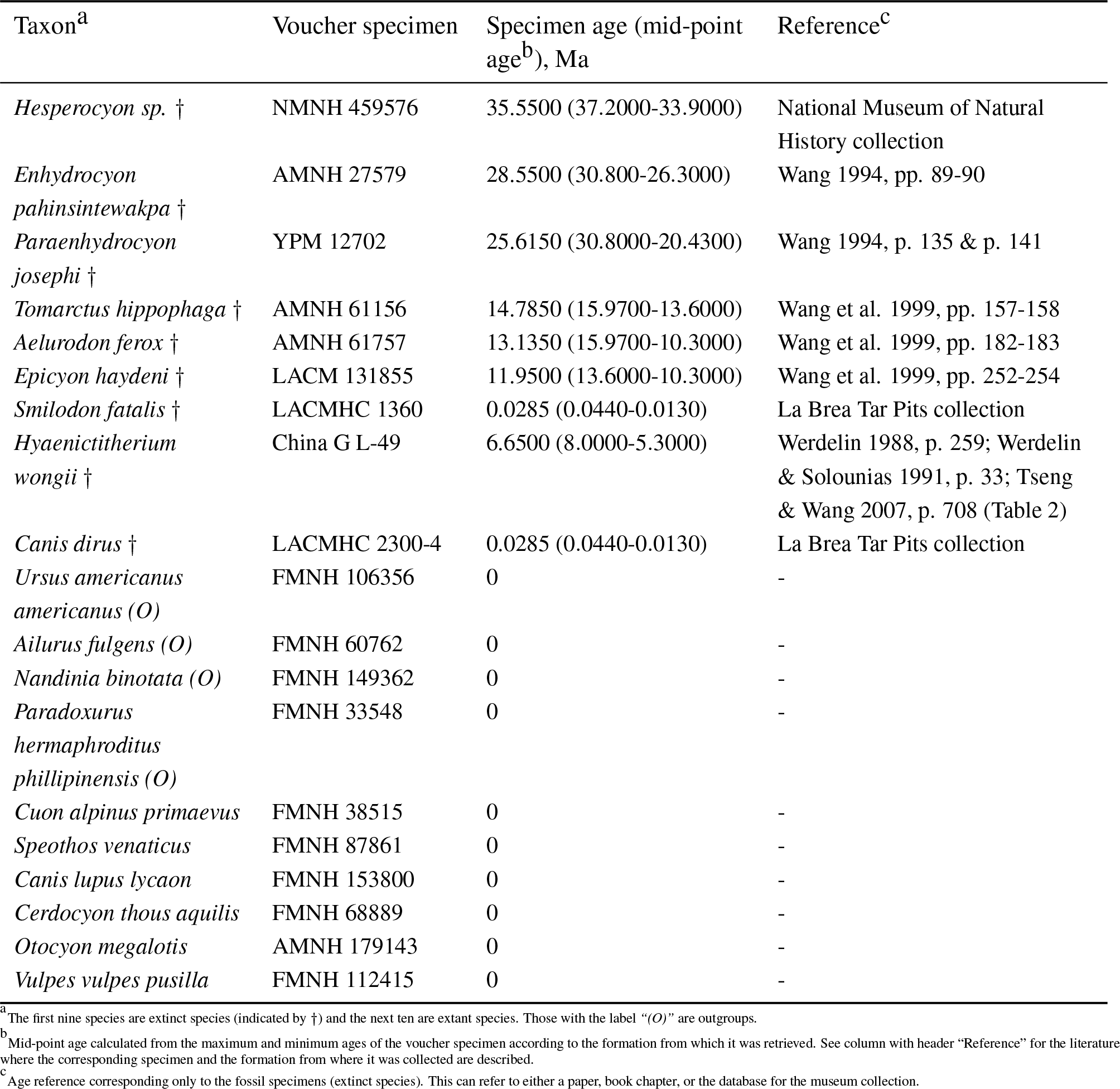
Summary of the 19 carnivoran species studied in this analysis. This table includes the voucher specimen, the specimen age and age ranges, and the reference for the specimen age and the age ranges. Note that, for the extant species, the specimen age is set to 0 as it refers to the present time.

**Figure 3:**
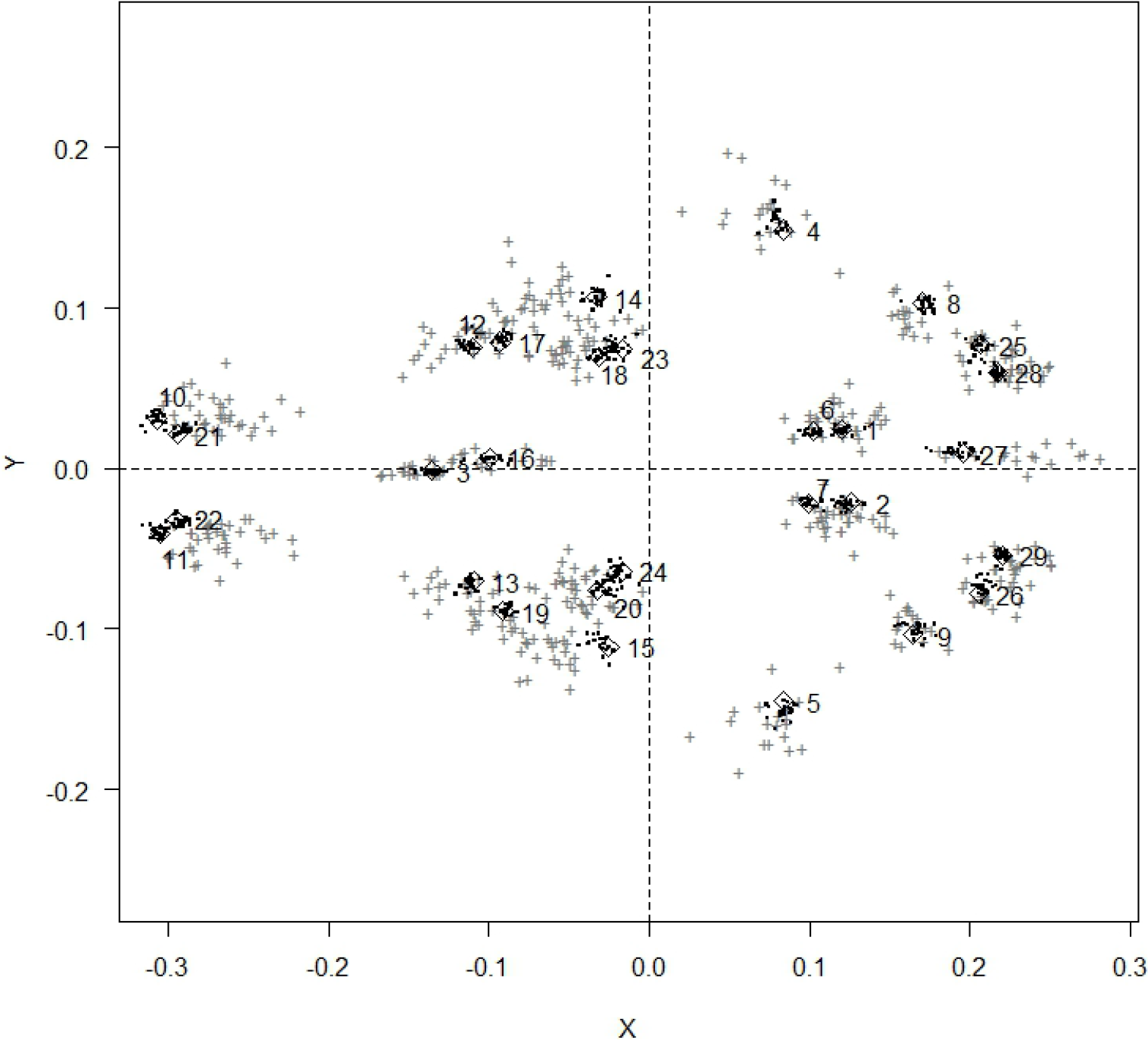
Procrustes alignment of 29 cranium landmarks for 19 carnivoran species. The alignment was obtained with the Morpho package in R. Landmark coordinates for 21 foxes (*Vulpes vulpes*) and 18 other carnivoran species are shown as dark grey crosses and black dots, respectively. The mean of the landmark coordinates are shown as diamonds and are numbered: 1, 2, Basioccipital-Basisphenoid-Bulla suture - (left, right); 3, Palatine - Maxilla - ventral suture; 4, 5, Jugal - Squamosal ventral suture (left, right); 6, 7, Bulla - anterior extreme - (left, right); 8, 9, Bulla - posterior lateral extreme - (left, right); 10, 11, Premaxilla - anterior extreme - (left, right); 12, 13, Jugal-Maxilla (Orbit crest) suture - (left, right); 14, 15, Jugal-Maxilla (base of zygomatic arch) suture - (left, right); 16, Nasals Frontal suture; 17, 19, Anterior lateral M1 - (left, right); 18, 20, Posterior lateral M2 - (left, right); 21, 22, Canine - mesial extreme - (left, right); 23, 24, Postorbital process tip - (left, right); 25, 26, Paraoccipital process tip - (left, right); 27, Parietals - Occipital suture; and 28, 29, Occipital condyle extreme - (left, right).

Landmark data are aligned using Procrustes superimposition (Gower, 1975; Rohlf and Slice, 1990), a technique in which the landmark coordinates for each individual are translated, rotated, and scaled to unit centroid size so the square of the distance between the individual’s landmarks and the mean landmark coordinates among all the individuals is minimised (see cited literature for details). We perform the Procrustes alignment in two steps. First, we align the 19 carnivoran species (excluding all but one of the foxes) using the `Morpho :: procSym` function in R (Schlager, 2017), resulting in a 19 × 87 geometric morphometric alignment **M**. Then, the remaining 20 foxes are aligned to the mean shape of **M** using the `Morpho :: align2procSym` function. This is done so that the alignment is not biased due to the large number of foxes. The resulting Procrustes alignment for the foxes, **P**(of size 21 × 87), is used to obtain the estimates of the population variances, **ĉ**, and the shrinkage correlation matrix, **R**\*, for the landmark coordinates. The correlation matrix **R**\* depends on the orientation of the landmarks, that is, different rotations of **P** may lead to different estimates of **R**\*. Therefore, **R**\* must be estimated on a population matrix that has been aligned to the species matrix. Divergence times are estimated on **Z**^(*s*)^, the transformed alignment obtained after scaling **M** by the population variances, and multiplying by the Cholesky decomposition of **R**\*. A summary of the methodology to generate the morphological alignment is given in Figure 4.

**Figure 4:**
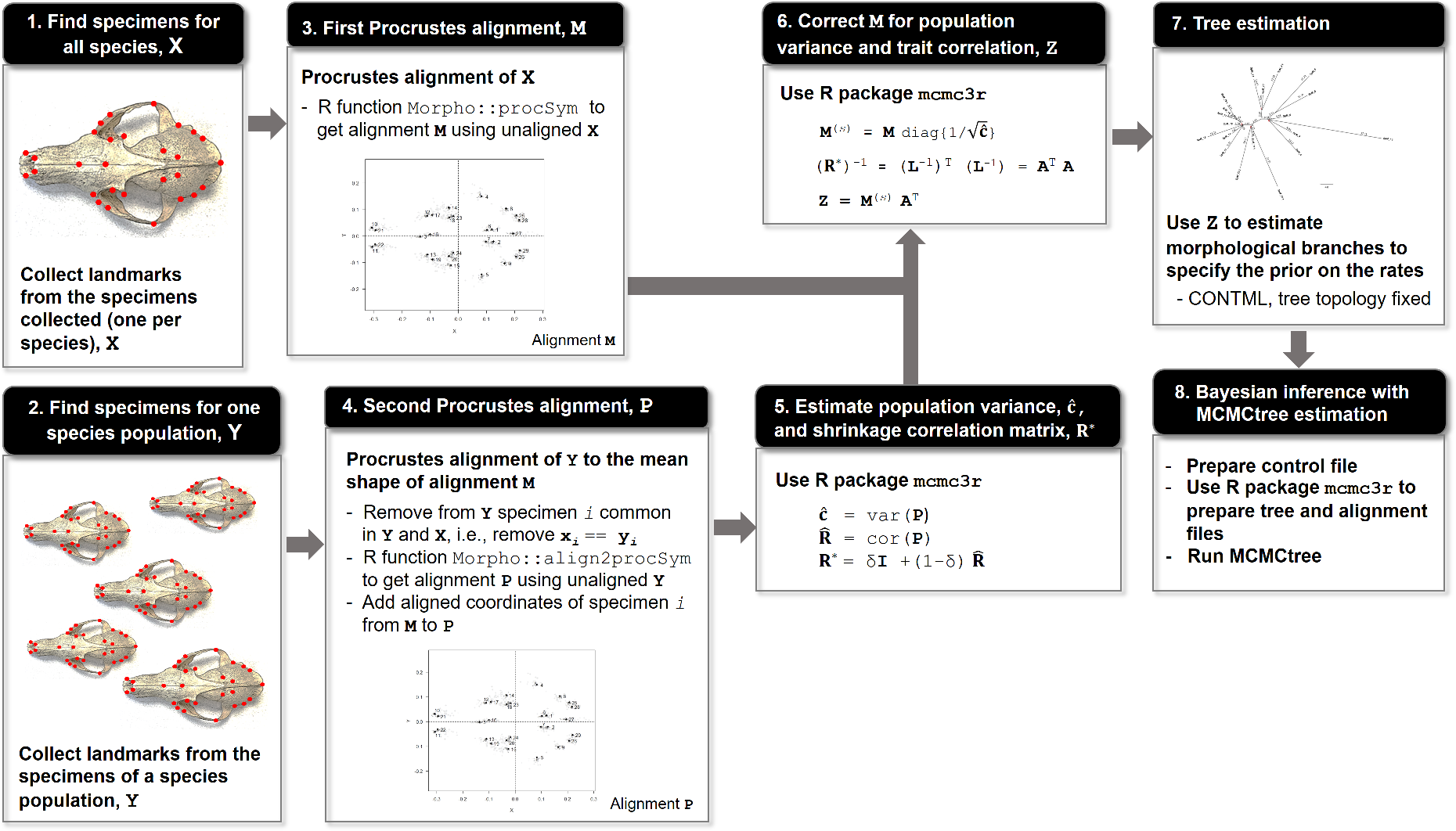
Summary of Bayesian inference with continuous landmark data. Step 1: Collect landmarks from the bones of the extinct and extant species and obtain matrix **X**. Step 2: Collect landmarks from the bones of a population sample of one of the species sampled in step 1 and obtain matrix **Y**. Step 3: Align the landmarks in **X** using the Procrustes method (for example using `Morpho :: procSym` in R) to obtain aligned matrix **M**. Step 4: Align landmarks from population sample in matrix **Y** to mean shape of alignment **M**(for example, with Morpho :: align2procSym) and obtain aligned population matrix **P**. Step 5: Use **P** to estimate population variance, **ĉ**, and shrinkage correlation matrix **R**\*. Step 6: Use **ĉ** to correct **M** for population noise and **R**\* to correct for within-lineage correlation among characters. This gives the corrected alignment **Z**. Step 8: Use **Z** in CONTML to estimate the morphological branches using a fixed tree topology (species tree). They are used to estimate the morphological rate and decide on the prior on rates. Step 8: Use the program MCMCtree to estimate divergence times and morphological rates of evolution. The `mcmc3r` package in R can be used to prepare the morphological alignment (i.e., to correct for within-lineage correlation and noise) and to generate the appropriate control files for MCMCtree.

### Molecular Data

We use the sequences of the 12 mitochondrial genes (mt-genes) for the 10 extant carnivoran species that are available at the NCBI: cytochrome c oxidase (*COX*) subunits 1, 2, and 3; cytochrome b (*CYTB*); NADH dehydrogenase (*ND*) subunits 1, 2, 3, 4, 4L, and 5; and ATP synthase F0 (*ATP*) subunits 6 and 8. We do not include *ND6* in our analysis because it is not encoded on the same strand of the mitochondrial DNA (mt-DNA) like the other 12 mt-genes, and thus has very different nucleotide compositions. Note that not all the 12 mt-genes are available at the NCBI for the 10 extant species analysed. Thus, gaps are introduced in the molecular alignment when a gene is not available for a species. Prank v.150803 (Löytynoja and Goldman, 2005, 2008) is used to align the molecular sequences. The concatenated gene alignment is divided into two partitions: (i) first and second codon positions (12CP) and (ii) third codon positions (3CP).

### Divergence Times Estimation

We estimate the divergence times with MCMCtree on the fixed carnivoran topology of Finarelli and Goswami (2009) and Martín-Serra et al. (2014). We use three data sets: (i) morphological alignment, (ii) molecular alignment in two partitions (12CP + 3CP), and (iii) morphological and molecular alignments (12CP + 3CP) analysed together as three partitions. The molecular data are analysed using the HKY+Γ (Hasegawa et al., 1984, 1985) substitution model, while the Brownian diffusion model of quantitative character evolution (Felsenstein, 1973) is used for the morphological data.

The prior on the ages of the nodes is constructed using the birth-death (BD) process (Yang and Rannala, 2006), if only extant species are analysed, or the birth-death-sequential-sampling (BDSS) model (Stadler and Yang, 2013), if fossil species are included in the analysis. For the BDSS prior we use *λ*_BDSS_ = *μ*_BDSS_ = 1, *ρ*_BDSS_ = 0 and *ψ*_BDSS_ = 0.001; and for the BD prior we use *λ*_BD_ = *μ*_BD_ = 1 and *ρ*_BD_ = 0.1. We chose both set of parameters to obtain approximately uniform prior distributions on node ages. Both the BDSS and BD processes are conditioned on the age of the root. Thus, we set a uniform fossil calibration with soft bounds on the root age between 37.3 Ma and 66 Ma, following Benton et al. (2015). The time unit is set to 1 myr.

We use a gamma-Dirichlet prior (dos Reis et al., 2014) on the (molecular and/or morphological) rate with shape *α* = 2 and with the scale parameter *β* chosen so that the mean of the prior rate (given by *α/β*) is close to empirical estimates based on the morphological or molecular branch lengths on the phylogeny. In the gamma-Dirichlet prior, one specifies the prior mean on the overall (average) rate for all partitions, then a Dirichlet distribution is used to partition the total rate among the partitions (see dos Reis et al., 2014 for details). To specify the prior, we first estimated, by maximum likelihood, branch lengths with RAxML v8.2.10 (Stamatakis, 2014) for the molecular alignment, and with CONTML (PHYLIP package, Felsenstein, 1993) for the morphological alignment. The resulting unrooted trees where midpoint rooted, and then we calculated a rough approximation to the number of substitutions, or units of morphological drift, from the tips of the root, and divided these by 52 Ma, the (rounded) midpoint value of the root calibration. This gives a rough idea of the value of the mean rates for the molecular and morphological partitions. These empirical rate estimates are then used to calculate the mean rate for the gamma-Dirichlet prior. Note that the use of *α* = 2 leads to a very diffuse (large variance) prior on the rate. The chosen values of *β* for all the data sets are given in Table 2. The data are analysed under the strict clock (STR), the geometric Brownian diffusion (GBM, also known as autocorrelated-rates, Thorne et al., 1998; Yang and Rannala, 2006), and independent log-normal rate (ILN) models (Rannala and Yang, 2007; Lemey et al., 2010). The gamma-Dirichlet prior on 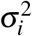 for the GBM and ILN models is 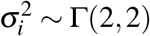 for both the molecular and the morphological data sets.

**Table 2:**
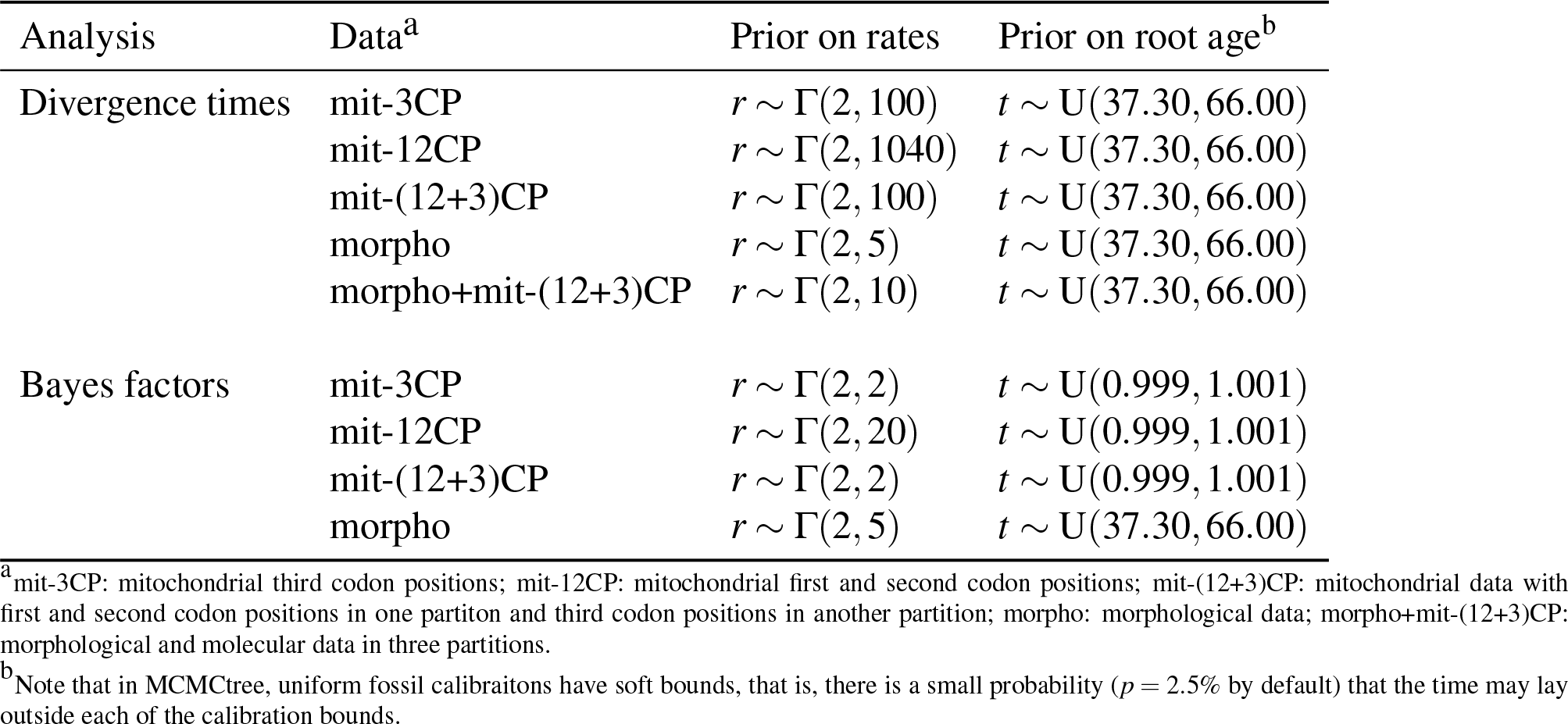
Priors on evolutionary rates and root age for the Carnivora analysis.

### Bayesian Selection of Clock and Correlation Models

We use Bayes factors (BFs) to select among the three clock models for the morphological and molecular data sets. Marginal likelihoods for each model are calculated using the stepping-stone approach (Xie et al., 2011) as implemented in the `mcmc3r R` package (dos Reis et al., 2018). The estimated marginal likelihoods are then used to calculate the BFs and posterior probabilities for each clock model. Note that when molecular data only were analysed, the age of the root is fixed to 1 (as there are no fossil tip species to calibrate the tree). In MCMCtree, this is done by using a narrow uniform distribution with soft bounds on the age of the root, U(0.999, 1.001). In this case, the mean of the rate prior needs to be modified to accommodate the different age of the root. Table 2 gives the modified priors.

Bayes factors can also be used to select for the correlation model in the morphological data. The marginal likelihood can be calculated by using **R** = **I** in Eq. (5), that is, by assuming characters evolve independently, or calculated on **Z**^(*s*)^ which has been transformed to account for the correlation among characters. Please note that, when using **Z**^(*s*)^, the likelihood of Eq. (5) must be scaled by the determinant |**R**\**|* so that the marginal likelihood is calculated correctly. The marginal likelihoods can then be used to calculate the BF and posterior probability for the independent and correlated models.

## Results

### Analysis of Simulated Data

In general, the simulation results met our expectations. We found that estimates of divergence times and rates for large number of characters and with older fossils were close to the true values. On the other hand, when the data sets were simulated with population noise and/or with correlated characters, but these were not corrected for, the estimated parameters were far from the true values. This bias was particularly large when the population variance was large or when the correlation among characters was very strong. We describe the results in detail below.

#### Effect of the number of characters and fossil age

Figure 5 shows the effect of sample size and fossil age on posterior estimates of the root age, *t*_9_, and morphological rate, *r*. Posterior means and 95% quantiles of *t*_9_ and *r* are averaged across all 1,000 simulation replicates and plotted. As expected, uncertainty (as measured by the CI width) in the estimates decreases for larger data sets and when the age of fossil H is the oldest. For example, when *t*_*H*_ = 0.1 and *p* = 100 characters are analysed, the average CI of *t*_9_ is 0.8-1.2, which is 0.4 time units wide, or 40% of the root age (Fig. 5A). However, this uncertainty is reduced to only 13% of the root age when analysing *p* = 10, 000 characters (Fig. 5C). The uncertainty is reduced even further when *t*_*H*_ = 0.7 (when the fossil is the oldest), giving a CI width which is about 5% of the root age estimate (Fig. 5C). Note that the younger the fossil is, the larger the distance from the fossil to the root of the tree is, which makes the fossil less informative. The same pattern is observed for the estimates of the morphological rate (Fig. 5A’-C’) and for the rest of the node ages (Tables S1 and S2). Note that, in all cases, the estimates appear unbiased and converging to the true values as the data become more informative.

**Figure 5:**
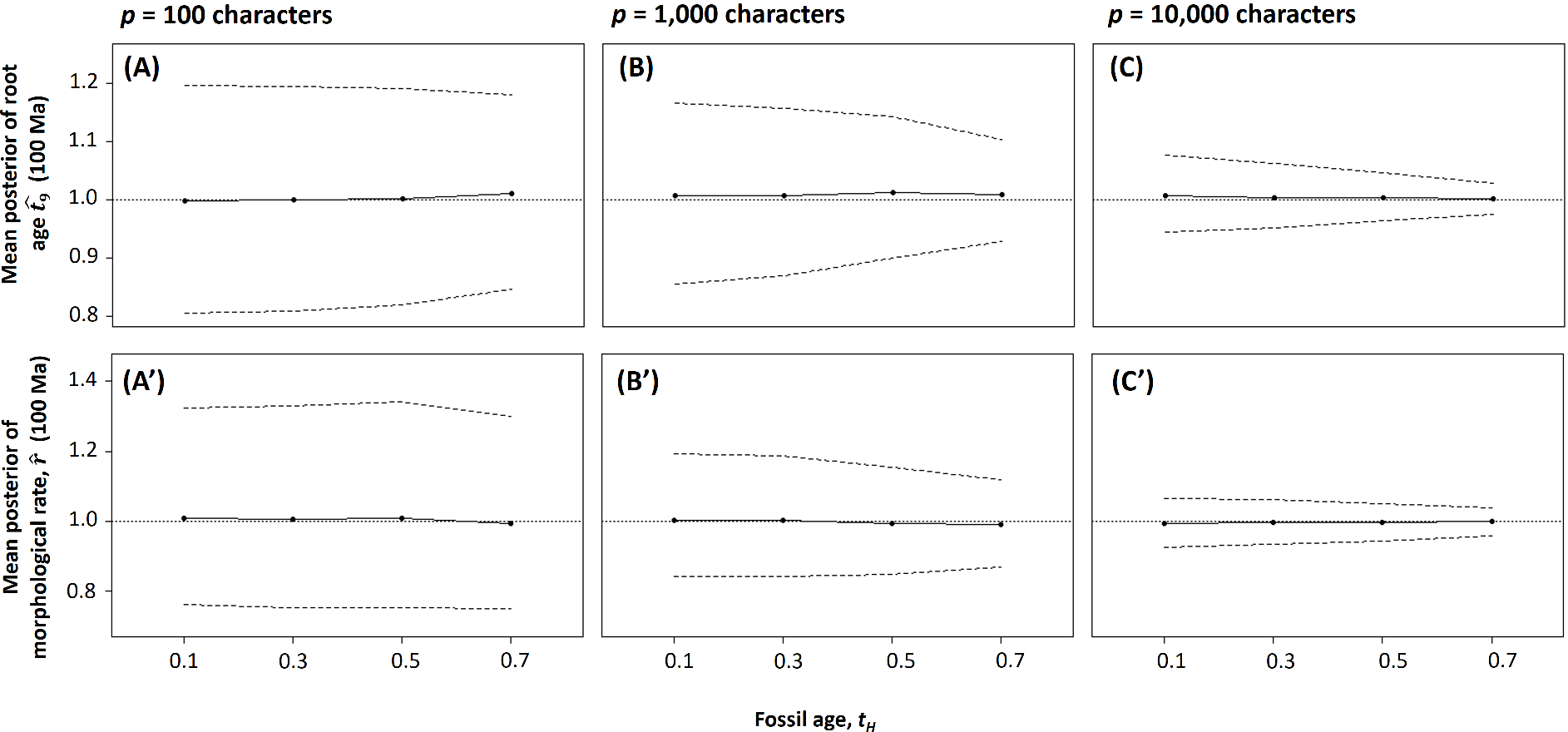
Effect of the number of characters and fossil age on posterior estimates of the root age and morphological rate for simulated morphological characters. The posterior mean and 95% quantile estimates of *t*_9_ and *r* are averaged over *R* = 1, 000 replicates. Quantitative characters were simulated under the phylogeny of Figure 2, and the age of fossil H, *t*_*H*_, was varied to study the effect of the fossil age on the estimates. The true root age, *t*_9_ = 1.0, and the true morphological rate, *r* = 1.0, are represented as horizontal dotted lines. The dashed lines give the corresponding upper and lower 95% CI limits.

#### Effect of population noise

Figure 6 shows the effect of the population noise on the estimates of the root age, *t*_9_, and morphological rate, *r*, when *p* = 1, 000 characters are analysed. As above, estimates are averaged across the 1, 000 replicates and plotted. When the population noise is ignored in the analysis (Fig. 6A and A’), the parameters are overestimated and the overestimation is largest for the largest population noise. For example, when *c* = 0.5 and when *c* is ignored in the analysis, the average of the posterior mean of *t*_9_ is 1.2 (Fig. 6A), which has a mean bias of *b* = 0.2 or a relative bias of 20%. This is a large bias that cannot be corrected by sampling more characters because the model is misspecified. On the other hand, when *c* = 0.5 and when **ĉ** is used to correct for the population noise in the analysis, the relative bias in the estimate of *t*_9_ is only about 4% (Fig. 6B). Note that we expect some bias to remain in the estimates because **ĉ** itself has sampling errors: we need to estimate one variance for each character, and these variance estimates are obtained from a small population sample of 20 individuals. Asymptotically, as the population sample increases to infinity, the sampling errors go to zero and **ĉ** would converge to the true population variances, **c**. In this case, we expect to see no bias in the posterior means of *t* and *r*. This is exemplified in Figure 6C, where the data has been scaled by the true variances, **c**, and thus there is almost no bias in the posterior mean of the root age. The pattern of bias in the estimates of *t*_9_ when the population noise is ignored in the analysis is also seen for estimates of the morphological rate, *r,* (Fig. 6A’-C’) and for the rest of the node ages in the phylogeny (Tables S3 and S4).

**Figure 6:**
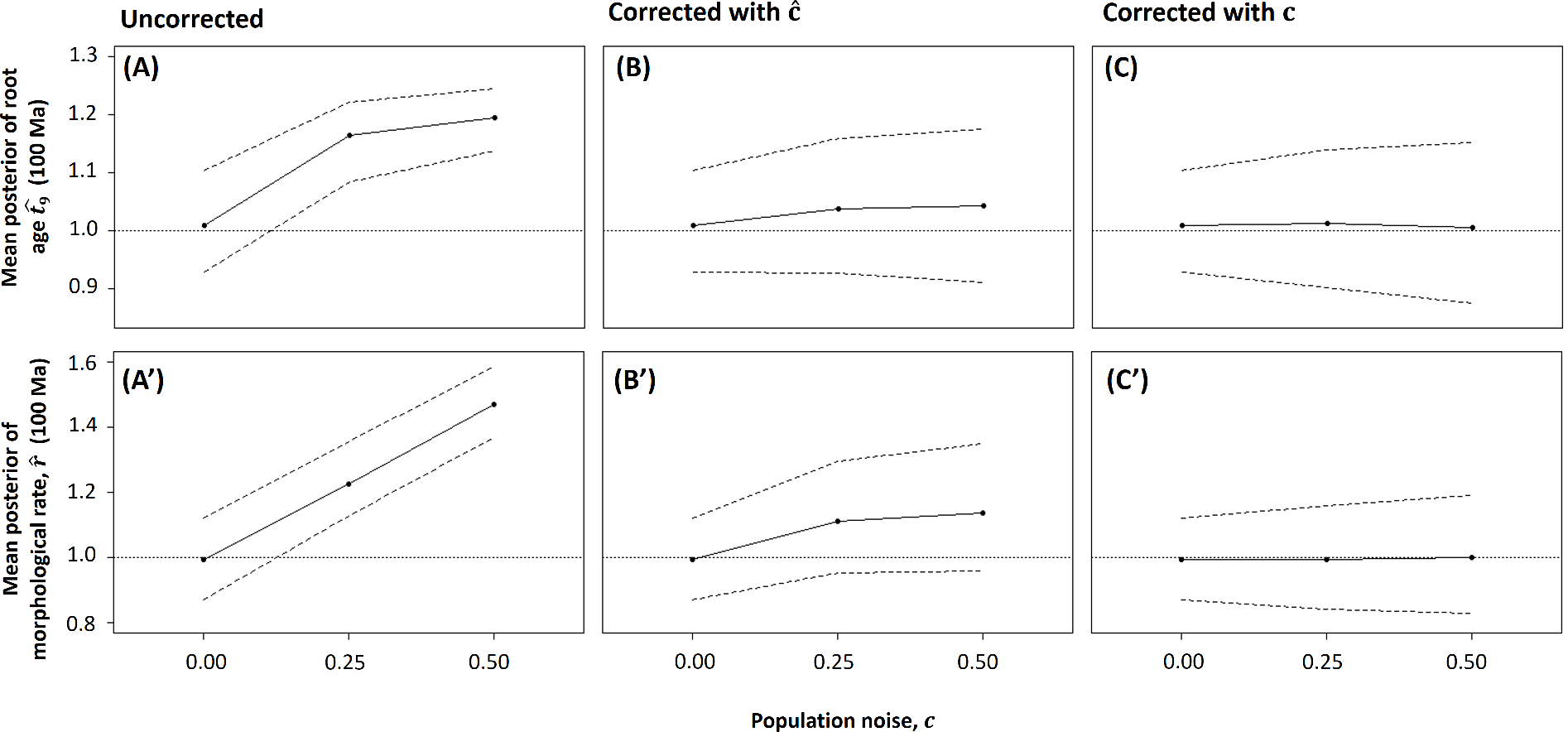
Effect of population noise on estimates of the age of the root and the morphological rate for simulated morphological characters. The posterior mean and 95% quantile estimates of *t*_9_ and *r* are averaged over the *R* = 1, 000 replicates. The *p* = 1, 000 quantitative characters were simulated under the phylogeny of Figure 2. (A, A’): the population noise is ignored during Bayesian inference, (B, B’): the population noise is corrected using the vector of estimated population variances, **ĉ**; (C, C’): the population noise is corrected using the vector of true population variances, **c**. The true root age, *t*_9_ = 1.0, and the true morphological rate, *r* = 1.0, are represented as horizontal dotted lines. The dashed lines give the corresponding upper and lower 95% CI limits.

#### Effect of correlation among characters

Figure 7 shows the effect of character correlation on estimates of the root age, *t*_9_, and the morphological rate, *r*, when *p* = 1, 000 characters are analysed and when the population noise is *c* = 0.25. As above, estimates are averaged across the 1,000 replicates and plotted. When both the population noise and the character correlation are ignored in the analysis, the time estimates tend to be more overestimated as the character correlation increases (Fig. 7A). For example, when *ρ* = 0.9 and when both correlation and noise are ignored, the average estimate of *t*_9_ = 1.42, with a bias *b* = 0.42 or relative bias of 42% (Fig. 7A). This is a very high bias in the estimate. Note that when *ρ* = 0.9 and the data are corrected for the population noise but not for the correlation, the large bias in the estimate of *t*_9_ remains (Fig. 7B). On the other hand, when *ρ* = 0.9 and both the noise and correlation are taken into account in the analysis, the bias in the estimate of *t*_9_ is very small (about 4%, Fig. 7C). This trend, in which *t*_9_ is overestimated when the character correlation is ignored, is also observed for the estimates of the other node ages (Tables S5 and S6).

**Figure 7:**
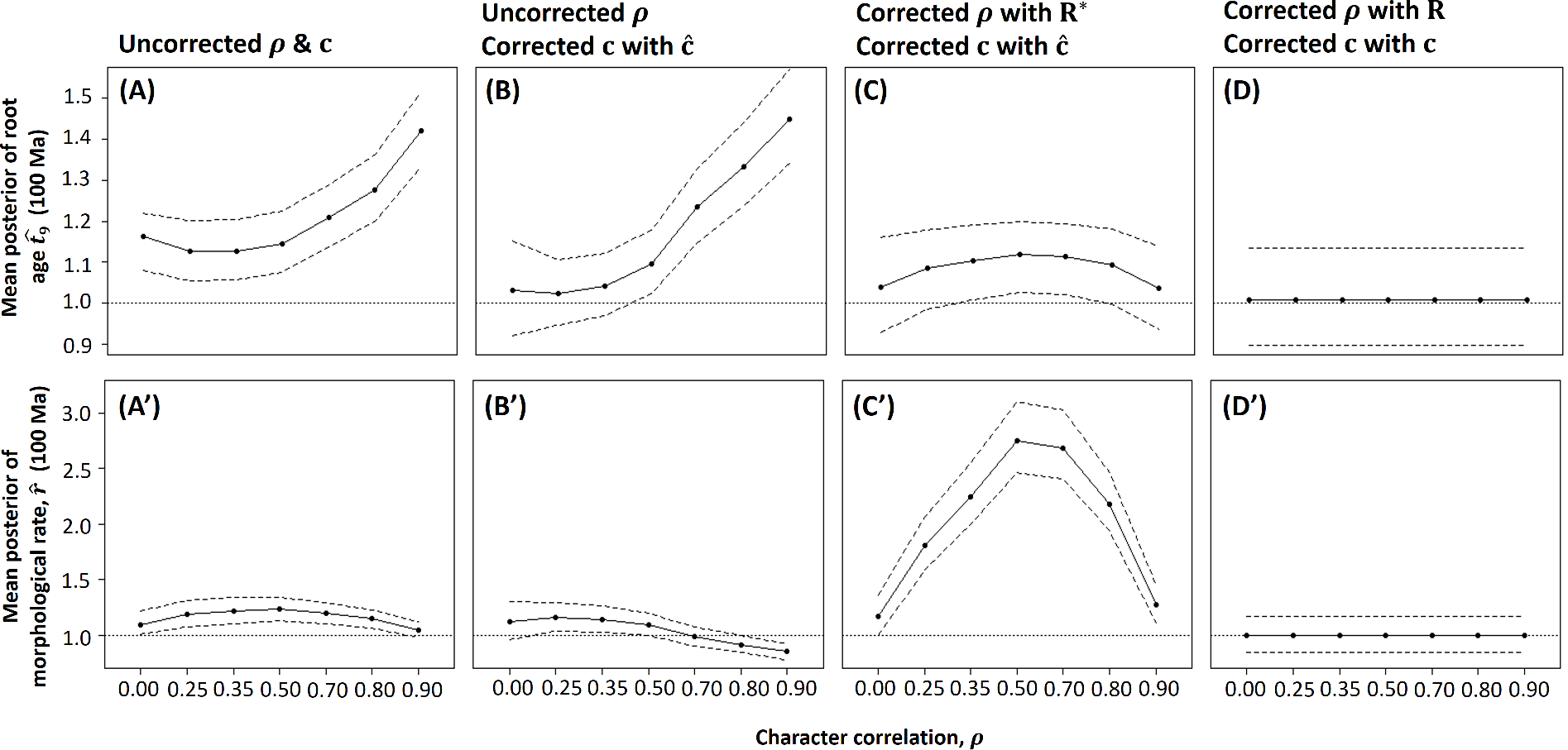
Effect of within-lineage correlation among characters on estimates of the root age and the morphological rate for simulated morphological characters. The posterior mean and 95% quantile estimates of *t*_9_ and *r* are averaged over the *R* = 1, 000 replicates. The *p* = 1, 000 quantitative characters were simulated under the phylogeny of Figure 2 with population noise *c* = 0.25. (A, A’): both population noise and within-lineage character correlation were ignored during Bayesian inference, (B, B’): within-lineage character correlation was not corrected for in the data sets but population noise was accounted for, (C, C’): both population noise and within-lineage character correlation were corrected for in the data sets, (D,D’): both population noise and within-lineage character correlation were corrected for the true values in the data sets. The true root age, *t*_9_ = 1.0, and the true morphological rate, *r* = 1.0, are represented as horizontal dotted lines. The dashed lines give the corresponding upper and lower 95% CI limits. Note that due to the strange pattern in C’, we extended the simulation analysis to include additional correlation values: *ρ* = 0.25, 0.35, 0.7, and 0.8.

Strangely, a different pattern is observed for the estimate of the rate. When the population noise and the character correlation are ignored in the analysis, or when the noise alone is corrected for, the bias in the estimate of *r* are moderate or small (Fig. 7A’-B’). Surprisingly, when *ρ* = 0.5 and when both the noise and character correlation are corrected for in the analysis, we find that the bias in the estimate of *r* is very high, an overestimation (relative bias) of about 175% (Fig. 7C’). The bias then decays to about 27% when *ρ* = 0.9 (Fig. 7C’). We note that these estimates are obtained when using the shrinkage estimate, **R**\*, to correct for the correlation. When using the unbiased estimate, **R̂**, to correct for the correlation, the errors in the estimates of the rate are so large that they cannot be included in Figure 7 (but see Tables S5 and S6). We note that both the estimates **R**\* and **R̂** are expected to contain large errors as we are estimating too many correlations from a small population sample. For example, when *p* = 1,000 characters we have to estimate 499,500 correlations. It appears that estimates of the morphological rate may be sensitive to errors in these estimates.

### Analysis of the Carnivora Data

#### Morphological tree and Smilodon landmarks

The morphological tree estimated with CONTML (PHYLIP package, Felsenstein, 1993) is shown in Figure S5. Because the branch length from the root of the tree to the extinct saber-tooth tiger, *Smilodon fatalis*, is very long, we examined the landmarks of this specimen for possible problems before Bayesian inference of divergence times. We used the function `geomorph :: plotOutliers` (Adams and Otárola-Castillo, 2013) in R to calculate the Procrustes distance from each specimen to the mean shape. The resulting plot (Fig. S3A) shows *Smilodon* as an outlier. In order to elucidate which landmarks place *Smilodon* as an outlier, we carried out a principal components analyses (PCA) of shape variation, with the first two components shown in Figure S4. Convex hull polygons were added to cluster the specimens: (i) Caniformia or Feliformia suborder, (ii) extant or extinct specimens, and (iii) outgroup or non-outgroup specimens. Moving along PC1 correlates with shrinking of the length of the cranium from the occipital to the maxillar, while PC2 correlates with an increase in the width of the cranium (Fig. S4). *Smilodon* is located at the extremes of both PCs, that is, it has an unusually short snout and a wide cranium. In other words, while all our specimens except *Smilodon* have dog- or bear-like skulls, *Smilodon* has a markedly different, emphatically cat-like shape. This explains the long branch for *Smilodon* in the morphological tree. Furthermore, *Smilodon* species have been found to be outliers in larger data sets too (Goswami et al., 2011). We keep *Smilodon* in the Bayesian analysis to illustrate the large variations in morphological rate in this phylogeny.

#### Bayesian selection of clock and correlation models

Table 3 shows the results of the Bayesian model selection. For the molecular data, the ILN rates model is best (*P* = 0.75) when the two molecular partitions are analysed jointly. However, when they are analysed separately, the GBM rates model is best for the third codon positions (*P* = 0.74), while the ILN is marginaly better for the first and second codon positions (*P* = 0.53). For the morphological data, the ILN rates model with character correlation is best (*P* = 1.00). It is worth noting that including character correlation in the model improves the marginal likelihood by over 100 likelihood units compared to the no-correlation model (that is, all clock models are over 100 likelihood units higher when including the correlation). In contrast, when accounting for correlation, the ILN model is only 12.38 and 73.55 likelihood units better than the GBM and STR rate models, respectively. This large likelihood increase for the correlation model emphasizes that correlation is an important feature of morphological data that should be taken into account in the analysis.

**Table 3:** Bayesian selection of clock and correlation model for the Carnivora data.

| Data<sup>a</sup> | Model<sup>b</sup> | $\log L \pm \text{S.E}$ | Pr |
| --- | --- | --- | --- |
| mit-3CP | **GBM** | **$-22,011.37 \pm 0.05$** | **0.74** |
| | ILN | $-22,012.41 \pm 0.05$ | 0.26 |
| | STR | $-22,019.57 \pm 0.04$ | 0.00 |
| mit-12CP | GBM | $-25,651.40 \pm 0.04$ | 0.47 |
|  | **ILN** | **$-25,651.28 \pm 0.04$** | **0.53** |
| | STR | $-25,657.82 \pm 0.03$ | 0.00 |
| mit-(12+3)CP | GBM | $-47,658.83 \pm 0.05$ | 0.24 |
|  | **ILN** | **$-47,657.71 \pm 0.05$** | **0.75** |
| | STR | $-47,694.37 \pm 0.03$ | 0.00 |
| Morpho | GBM - ($\mathbf{R} = \mathbf{R}^*$) | $-4,097.41 \pm 0.04$ | 0.00 |
| | GBM - ($\mathbf{R} = \mathbf{I}$) | $-4,221.13 \pm 0.04$ | 0.00 |
|  | **ILN - ($\mathbf{R} = \mathbf{R}^*$)** | **$-4,085.03 \pm 0.02$** | **1.00** |
| | ILN - ($\mathbf{R} = \mathbf{I}$) | $-4,207.59 \pm 0.02$ | 0.00 |
| | STR - ($\mathbf{R} = \mathbf{R}^*$) | $-4,158.38 \pm 0.01$ | 0.00 |
| | STR - ($\mathbf{R} = \mathbf{I}$) | $-4,280.18 \pm 0.01$ | 0.00 |

<sup>a</sup>mit-12CP: 1 partition with the first and second codon positions (12CP) of the 12 concatenated mitochondrial genes (12-mit genes); mit-3CP: 1 partition with the third codon positions (3CP) of the 12-mit genes; mit-(12+3)CP: the two mitochondrial partitions analysed jointly; Morpho: 1 partition with the morphological alignment of 87 characters for the carnivoran data set.
<sup>b</sup>STR: strict clock model, GBM: autocorrelated-rates model, ILN: independent-rates model, $\mathbf{R} = \mathbf{I}$ : no correlation model (i.e., $\mathbf{R} = \mathbf{I}$ in Eq. 5), $\mathbf{R} = \mathbf{R}^*$ : correlation model (i.e., $\mathbf{R} = \mathbf{R}^*$ in Eq. 5). Note that, in all cases, $c = 1$ , that is, population noise is explicitly accounted for in the models.

#### Divergence time estimation

All divergence time estimates are obtained under the ILN rates model. Figure 8 shows the time calibrated Carnivora phylogeny. Posterior estimates using molecule-only (Fig. 8E), morphology-only (Fig. 8A,C,D), and joint (molecule and morphology, Fig. 8B) data sets are consistent with each other as the 95% HPDs of all analyses overlap. However, for some nodes in the phylogeny (e.g., the *Canis-Vulpes* extant clade), estimated dates are younger for the molecular data. Interestingly, the most precise estimates (i.e., with the narrowest HPDs) are obtained from the joint analysis of morphological and molecular data. Table 4 gives a summary of posterior estimates for the age of the root and extant clade *Canis-Vulpes* as well as the morphological and molecular rates. Our estimates for the *Canis-Vulpes* divergence time, which roughly vary between 13–37 Ma (depending on analysis), overlap with the estimates (23–38 Ma) of dos Reis et al., 2012. However, our results are in general older than those of Matzke and Wright, 2016, who report several analyses of discrete morphological characters for various canids. They gave their best estimates for Caninae divergence to be around 10 Ma (but as old as 40 Ma for unrealistic analyses settings).

**Table 4:** Posterior estimates of times (root and canid nodes) and rates for the Carnivora data under the ILN rates model.

| Model<sup>a</sup> | Time estimates (95% HPD interval)<sup>b</sup> | Rate estimates (95% HPD interval)<sup>c</sup> | Log-normal shape parameter (95% HPD interval) | Coefficient of rate variation<sup>d</sup> |
| --- | --- | --- | --- | --- |
| **Morphological data**<br>($\mathbf{R} = \mathbf{R}^*, \mathbf{c} = 1$) | $t_{\text{root}} = 54.6$ (42.7, 65.8)<br>$t_{\text{canid}} = 23.8$ (13.2, 36.2) | $\bar{r}_{\text{morpho}} = 0.488$ (0.284, 0.838) | $\sigma^2_{\text{morpho}} = 1.15$ (0.540, 2.10) | $\text{CV}_{\text{morpho}} = 1.47$ |
| **Morphological data**<br>($\mathbf{R} = \mathbf{I}, \mathbf{c} = 1$) | $t_{\text{root}} = 52.4$ (41.2, 65.3)<br>$t_{\text{canid}} = 25.4$ (14.6, 37.4) | $\bar{r}_{\text{morpho}} = 0.492$ (0.287, 0.844) | $\sigma^2_{\text{morpho}} = 1.10$ (0.485, 2.10) | $\text{CV}_{\text{morpho}} = 1.42$ |
| **Morphological data**<br>($\mathbf{R} = \mathbf{I}, \mathbf{c} = 0$) | $t_{\text{root}} = 52.1$ (41.5, 64.9)<br>$t_{\text{canid}} = 26.3$ (15.5, 38.1) | $\bar{r}_{\text{morpho}} = 0.491$ (0.288, 0.849) | $\sigma^2_{\text{morpho}} = 1.10$ (0.482, 2.08) | $\text{CV}_{\text{morpho}} = 1.42$ |
| **Molecular data** | $t_{\text{root}} = 45.5$ (36.4, 63.5)<br>$t_{\text{canid}} = 21.7$ (15.3, 31.7) | $\bar{r}_{\text{mit12}} = 0.0044$ (0.0028, 0.0065)<br>$\bar{r}_{\text{mit3}} = 0.0319$ (0.0207, 0.0451) | $\sigma^2_{\text{mit12}} = 0.1673$ (0.0353, 0.483)<br>$\sigma^2_{\text{mit3}} = 0.1131$ (0.0262, 0.321) | $\text{CV}_{\text{mit12}} = 0.43$<br>$\text{CV}_{\text{mit3}} = 0.35$ |
| **Joint (Molecular and Morphological),**<br>$\mathbf{R} = \mathbf{R}^*, \mathbf{c} = 1$) | $t_{\text{root}} = 52.0$ (41.7, 64.6)<br>$t_{\text{canid}} = 25.1$ (18.7, 32.7) | $\bar{r}_{\text{morpho}} = 0.452$ (0.268, 0.766)<br>$\bar{r}_{\text{mit12}} = 0.0037$ (0.0026, 0.0052)<br>$\bar{r}_{\text{mit3}} = 0.0273$ (0.0193, 0.0382) | $\sigma^2_{\text{morpho}} = 1.017$ (0.468, 1.94)<br>$\sigma^2_{\text{mit12}} = 0.159$ (0.0326, 0.456)<br>$\sigma^2_{\text{mit3}} = 0.147$ (0.0320, 0.425) | $\text{CV}_{\text{morpho}} = 1.33$<br>$\text{CV}_{\text{mit12}} = 0.42$<br>$\text{CV}_{\text{mit3}} = 0.40$ |

<sup>a</sup> $\mathbf{R} = \mathbf{R}^*$ : means the shrinkage estimate of the correlation matrix is used. $\mathbf{R} = \mathbf{I}$ : means the correlations are ignored, that is, the data are assumed to be independent and the correlation matrix is the identity matrix. $\mathbf{c} = 1$ and $\mathbf{c} = 0$ : means the population noise is corrected for or ignored, respectively, in the analysis.
<sup>b</sup> $t_{\text{canid}}$ refers to the age of the divergence of the extant *Canis-Vulpes* group.
<sup>c</sup>Here, $\bar{r}$ refers to the posterior estimate of the mean of the rate among branches.
<sup>d</sup>The coefficient of variation of the log-normal distribution of rates is $\text{CV} = \sqrt{\exp(\sigma^2) - 1}$ .

**Figure 8:**
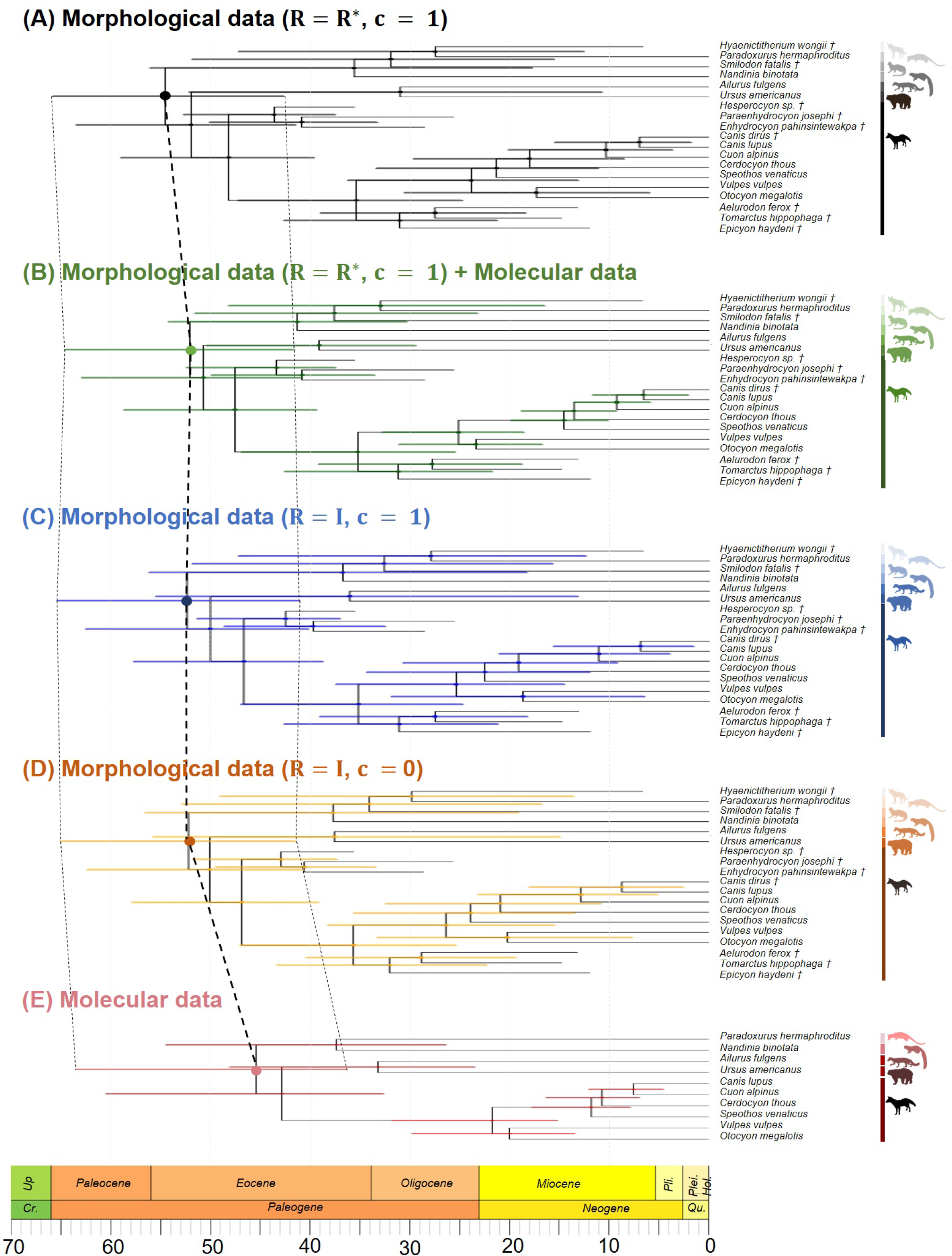
Divergence times for the 19 carnivoran species estimated with MCMCtree using morphological-only, molecular-only, and joint (morphological and molecular) data sets: (**A**) morphological-only data set accounting for population noise and within-lineage character correlation, (**B**) joint data set with the morphological data set in (A), (**C**) morphology-only data set without correcting for within-lineage character correlation and ignoring population noise despite having scaled the morphological matrix, (**D**) morphology-only data set without correcting for within-lineage character correlation nor population noise, and (**E**) molecule-only data set. Horizontal bars are the HPD of node ages. Calibration for the root: U(37.3, 66.0). ****Cr.****: Cretaceous, ***Up***.: Upper/Late, ***Pli.***: Pliocene, ***Plei.***: Pleistocene, ***Hol.***: Holocene, ***Qu.***: Quaternary. The posterior estimates for the root age (*t*_*root*_) and the corresponding 95% CIs are highlighted for each data set, the former connected through a bold dashed line and the latter through two corresponding dotted lines.

An interesting finding is that there is much more rate variation in the morphological rates than in molecular rates. In other words, molecular rates are more clock-like than morphological ones. For example, the coefficient of variation, 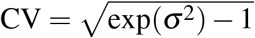, where *σ* ^2^ is the shape parameter (or log-variance) for the log-normal distribution, ranges between 1.3-1.8 for morphological characters and between 0.3-0.4 for the molecular data (Table 4). This indicates that morphological rates are three to four times more variable than molecular data.

Note that for the scaled landmark data, the within-population variances are set to *c* = 1. Under the ILN model, the estimated mean amount of morphological evolution from the root of the phylogeny to the tip is *r̅*_morpho_ × *t*_root_ = 0.49 × 52 = 25.5. Thus, the population variance represents 1/25.5 = 3.9% of the total expected morphological branch length from the root to the tip. That is the amount by which the external branches are extended due to the population noise. The estimated **ĉ** and **R**\* for the Carnivora data are given as Supplementary Material, and also given as example data in our `mcmc3r` package (which the user can use to reproduce the full Carnivora analysis presented here).

## Discussion

### Character Correlation

Our simulations highlight the importance of accounting for character correlation and population noise when continuous morphological data are used for divergence time estimation. However, when both factors are accounted for, we observed an unexpected result in our simulation study: the larger the correlation, the smaller the error to estimate both divergence times and evolutionary rate. Furthermore, the largest error occurred when *ρ* = 0.50, and the error was more dramatic on the rate estimates (see Fig. 7C and C’). The reasons for this are not clear to us, but we speculate that this may be due to the use of the shrinkage correlation matrix, **R**\*. Estimating the character correlations is a notoriously difficult task (e.g., Goolsby, 2016) as usually the number of characters is much larger than the number of samples, and thus the traditional estimate of the covariance matrix cannot be inverted. Therefore, it may be a worthwhile effort to assess the effects of different approaches to estimate the correlation matrix (e.g., Clavel et al., 2018). Other such approaches include matrix bending (e.g., Meyer and Kirkpatrick, 2010) or Bayesian estimation of the correlation matrix. The latter approach offers good prospects as the Bayesian estimate of the matrix would be regularised by the use of a prior, leading to well behaved estimates. The Wishart distribution (a multivariate generalisation of the gamma distribution) is the conjugate prior of the precision matrix (the inverse of the covariance matrix) and can thus be used to obtain the posterior of the precision matrix analytically from a population sample. From this posterior we could then obtain samples of the precision matrix during MCMC, and use them to obtain the data transformation (Eq. 6). This approach, although computationally expensive, has the advantage of incorporating the uncertainty about the correlation estimates into the analysis.

In this paper we assumed the correlations among characters are the same throughout the phylogeny. The model follows Felsenstein (1973), who suggested estimating the covariances among characters from population samples (from one or more species), and then using these to calculate the Mahalanobis distance among the populations. This distance can then be used in the likelihood calculation. Let **x** = **m**_*i*_ − **m**_*j*_ be the vector of differences among the characters in populations *i* and *j*. Then *D*^2^ = **x**^T^**x** is the square of the Euclidean distance between **m**_*i*_ and **m**_*j*_. If population samples are available, we may obtain the covariance estimate, **Ĉ**. The square of the Mahalanobis distance is then defined as *M*^2^ = **x**^T^**Ĉx**. Note that the exponent of the node likelihood (Eq. 9) is proportional to the Mahalanobis distance, thus by plugging the Mahalanobis distances into the likelihood calculation we can accommodate the covariance among characters (Felsenstein, 1973). Our approach here, using the transform 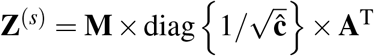, is equivalent to the Malahanobis method proposed by Felsenstein (1973), because *M*^2^ = **z**^(*s*)T^**z**^(*s*)^.

The assumption of constant correlations among lineages appears reasonable for closely related species, but may need to be relaxed when analysing more distantly related clades. For example, different covariance matrices can be estimated for different populations. Then the population-specific covariances could be used to calculate the likelihood for the terminal branches corresponding to the given populations. We could then use a stochastic process to model the changes in correlations across branches in the phylogeny and use this to sample the ancestral correlations using MCMC. However, this approach would be computationally very expensive. Revell and Harmon (2008) and Caetano and Harmon (2017) discuss further approaches to deal with variation of the correlation matrix along the phylogeny. In any case, assuming a constant correlation among lineages appears to be much better than assuming within-lineage independence among the characters. Here, for our Carnivora analysis, the best model with correlations is over 120 log-likelihood units better than the best independent model, and the posterior probability for the independent model is essentially zero (Table 3).

### Rate Variation Among Characters and Measurement Error

Felsenstein (1973) has shown that for a quantitative polygenic character with no dominance and under no selection, the rate of change for the character within a lineage is *r*_*k*_ ∝ *c*/*N*_*e,k*_, where *c* is the within-population variance of the character and *N*_*e,k*_ is the effective population size within the lineage. The population variance is 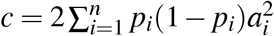, where *n* is the number of loci controlling the character, *p*_*i*_ and 1 − *p*_*i*_ are the allele frequencies at the (two-allele) *i*-th locus, and *a*_*i*_ is the contribution of each allele to the character value. Such a character will, asymptotically, be normally distributed as the number of loci increases (Fisher, 1919). Thus, different characters will have different within-population variances depending on the number of loci involved and the contribution of each loci to the value of the given character.

This among-characer variation can be modelled. However, this does not appear to be a worthwhile effort if character variances can be estimated from population samples. Let the relative rate of evolution for the *j*-th character be *g*_*j*_. Then, the length of the *k*-th branch in the phylogeny for the *j*-th character is *g*_*j*_*v*_*k*_ if the branch is an internal branch, and *g*_*j*_(*v*_*k*_ + *c*) if it is an external branch, where *g*_*j*_*c* is then the population variance for the character (which, as shown above, is proportional to the evolutionary rate). If we assume that the rates, *g*_*j*_, follow a discretised gamma distribution (or any other suitable distribution, e.g., Schraiber et al., 2013), then it is possible to integrate the among character rates out during calculation of the character likelihood as described in Yang (1994). However, because *g*_*j*_*c* (the character variance) can be estimated directly from a population sample and used to re-scale the characters, it turns out that the expectation of the re-scaled branch lengths is *g*_*j*_(*v*_*k*_ + *c*)/(*g_j_c*) = *v_k_/c* + 1 if the branch is an external branch, and *v_k_/c* if it is an internal branch. That is, the character rate, *g*_*j*_, drops out and the re-scaled branches are the same for all characters. Therefore, there is no need for a model of rate variation among characters. In practice, the estimates of the character variances contain sampling errors that will affect the asymptotic behaviour of the estimates (Fig. 6). Note that there is an important relationship between the among character rate variation and the within-lineage covariances of Eq. (9), thus we can always write 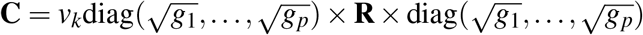.

The population variance of a trait will be similar across lineages if the number of loci is large or if the allele frequencies are similar across the populations (Felsenstein, 1973). However, if the number of alleles controlling the trait is small and if the allele frequencies are very different across populations, then *c* may vary among populations (Felsenstein, 1973). Let *c*^(*i*)^ be the population variance in species *i*. We can set *c*^(*i*)^ to be proportional to the morphological rate of the external branch for the given species (because *r*_*i*_ ∝ *c*^(*i*)^/*N*_*e,i*_). In this way, variation in *c* among species would become incorporated within the relaxed-clock model of rate variation among lineages. If a population sample for the *i*-th species is used to scale the characters to have unit variance, then we fix *c* ^(*i*)^ = 1 and set *c*^(*j*)^ to be proportional to the ratio 1/*r*_*i*_.

Quantitative characters may be subject to measurement errors (Ives et al., 2007). For example, landmark measurements may be subject to errors by the way a user identifies a landmark point, and landmark measurements may vary even when measured by the same user. In our carnivoran data, all specimens were measured by one of the co-authors. Thus, in our case, the measurement error is confounded with the population variance. This is unimportant as the confounded parameter is then used to correctly rescale the alignment for all characters. The effect of measurement error when measurements are obtained by different operators is a matter that will require further study and perhaps explicit modelling within our Bayesian framework (see Ives et al., 2007 for discussions).

### Limitations of the Brownian Diffusion Model

The Brownian diffusion model has a few undesirable features: the displacement (change) of a character is independent of its current state, there is no stationary distribution, and the variance in character change tends to infinity with time. These may be unrealistic for analysis of real data. For example, cranium landmarks are not expected to drift to arbitrarily large values for distantly related species. Alternative models include the Ornstein-Uhlenbeck model (OU, Lande, 1976; Hansen, 1997; Butler and King, 2004) or the Lévy processes (Landis et al., 2013). The former is an extension of the Brownian diffusion that stabilizes the displacement towards an optimum value (and thus has a stationary distribution and finite variance) while the latter is the sum of a directional drift, a Brownian diffusion, and a saltational jump in the character space. Parins-Fukuchi (2018b,a) has studied inference of phylogeny under the Brownian diffusion model for simulated and real data (including morphometric data for extant and extinct fossils) and found that the Brownian model performed well. Implementation of the OU model for Bayesian inference of topology and divergence times in a phylogeny appears worthwhile and a matter for future work.

### Partitioning the Morphological Alignment

The geometric morphometrics analyses carried out with the Carnivora data suggest that different partitioning schemes with morphological data sets should be explored. For instance, the results from the PCA (Fig. S4) indicate two regions within the carnivoran skulls that might follow different patterns of evolution: (i) from the maxillar to the lateral and (ii) from the lateral to the occipital. Previous research has shown different modules of correlated continuous characters are expected to evolve at different rates (Goswami et al., 2014; Felice and Goswami, 2018), suggesting the use of an appropriate partitioning scheme could improve the estimation of divergence times (Lee, 2016). Therefore, it would be interesting to explore the evolution of the cranium shape in this phylogeny when partitioning the data set into these two modules. Although this was not the aim of this study, we believe that partitioning morphological alignments according to modules identified using geometric morphometrics could improve estimates of rates and divergence times. This is particularly important for the morphological data because the evolutionary clock appears to be seriously violated, with some species showing very large rate variation (for example, *Smilodon*).

For example, Ho (2014) discusses how patterns of molecular rate variation may change for different regions of the genome. If these patterns of molecular rate variation are reflected on the morphological rates, then it may be worthwhile exploring whether partitioning morphological data would allow us to estimate these patterns. Methods for partitioning molecular data according to rate variation have been developed (Duchêne et al., 2014; Foster and Ho, 2017; Angelis et al., 2018), and these could in principle be combined with methods to detect morphological modules (partitions) based on morphological and/or developmental rates (e.g., Felice and Goswami, 2018). Note that if characters are scaled to have the same variance, then the overall rate for different character partitions will be the same. However, the *pattern* of rate variation among lineages (branches) and between partitions will be different. By incorporating morphological partitions with different patterns of rate variation among lineages, it should be possible to improve the precision of time estimates.

### Conclusions

The development of the total-evidence dating approach using discrete characters (Pyron, 2011; Ronquist et al., 2012) has allowed us to incorporate fossil data within an explicit modelling framework. Incorporation of continuous characters in the analysis is the natural extension of this framework. Recently, Parins-Fukuchi (2018b,a) used Felsenstein (1973) implementation of the Brownian model of character evolution to study in detail the performance of phylogenetic inference under the model on simulated and real data, assuming character independence and with emphasis on the ability of the model to place fossil taxa on the phylogeny. Our work here extends the Bayesian analysis of continuous characters by explicitly accounting for character correlation and population variance among the characters, and by the use of Bayesian selection of morphological rate model. Our results and those by Parins-Fukuchi (2018b,a) indicate the analysis of continuous characters is promising for the estimation of topology and divergence times in phylogenies. Perhaps the main advantage of using continuous characters is the easiness with which correlations can be incorporated in the analysis. In the Mk model, character correlation can be incorporated by expanding the model’s transition matrix to accommodate all the possible combinations of character transitions given the correlations (Pagel, 1994), with the resulting transition matrices becoming very large (Felsenstein, 2005). For example, to analyse *p* = 100 correlated binary characters, we would require a 2^100^ × 2^100^ transition matrix. The number of parameters to be estimated in this case, 8 × 10^59^, is larger than the number of atoms in the sun. In contrast, in the continuous case we would only need to estimate (*p*^2^ − *p*)/2 = 4, 950 correlations. Given that correlated character evolution is the rule rather than the exception, it appears that models that explicitly incorporate correlations are urgently required. The way forward appears to be the use of continuous characters, or the use of the threshold model for discrete characters, which explicitly incorporates a continuous process in the background (Felsenstein, 2005, 2012). If the discrete characters are ordered and can be assumed to have a continuous basis, then correlation can be introduced in the continuous variable (called liability), before it is discretized, as in the implementation of the auto-discrete-gamma model (Yang, 1995).

## Supplementary Material

Data available from the Dryad Digital Repository: http://dx.doi.org/10.5061/dryad/[NNNN].

## Acknowledgements

We would like to thank Jeff Thorne, Michael Landis, Simon Ho, Adam Leaché, Andrew Knapp and an anonymous reviewer for constructive comments and ideas. This study used Queen Mary’s Apocrita high-performance computer cluster (King et al., 2017).

## References

Adams, D. C. and E. Otárola-Castillo. 2013. geomorph: an R package for the collection and analysis of geometric morphometric shape data. Methods Ecol. Evol. 4:393–399.

Angelis, K., S. Álvarez-Carretero, M. Dos Reis, and Z. Yang. 2018. An evaluation of different partitioning strategies for Bayesian estimation of species divergence times. Syst. Biol. 67:61–77.

Arcila, D., R. A., Pyron, J. C., Tyler, G. Ortí, and R. Betancur-R. 2015. An evaluation of fossil tip-dating versus node-age calibrations in tetraodontiform fishes (Teleostei: Percomorphaceae). Mol. Phylogenet. Evol. 82:131–145.

Benton, M. J. and P. C. J. Donoghue. 2007. Paleontological evidence to date the tree of life. Mol. Biol. Evol. 24:889–891.

Benton, M. J., P. C. J. Donoghue, R. J., Asher, M. Friedman, T. J., Near, and J. Vinther. 2015. Constraints on the timescale of animal evolutionary history. Palaeontol. Electron. 18.1.1FC:1–106.

Butler, M. A. and A. A. King. 2004. Phylogenetic comparative analysis: a modeling approach for adaptive evolution. Am. Nat. 164:683–695.

Caetano, D. S. and L. J. Harmon. 2017. ratematrix: An R package for studying evolutionary integration among several traits on phylogenetic trees. Methods Ecol. Evol. 8:1920–1927.

Clavel, J., L. Aristide, and H. Morlon. 2018. A penalized likelihood framework for high-dimensional phylogenetic comparative methods and an application to New-World monkeys brain evolution. Syst. Biol. 0:1–25.

Donoghue, P. C. J. and M. J. Benton. 2007. Rocks and clocks: calibrating the Tree of Life using fossils and molecules. Trends. Ecol. Evol. 22:424–431.

dos Reis, M., P. C. J. Donoghue, and Z. Yang. 2016. Bayesian molecular clock dating of species divergences in the genomics era. Nat. Rev. Genet. 17:71–80.

dos Reis, M., G. F., Gunnell, J. Barba-Montoya, A. Wilkins, Z. Yang, and A. D. Yoder. 2018. Using phylogenomic data to explore the effects of relaxed clocks and calibration strategies on divergence time estimation: primates as a test case. Syst. Biol. 67:594–615.

dos Reis, M., J. Inoue, M. Hasegawa, R. J., Asher, P. C. J. Donoghue, and Z. Yang. 2012. Phylogenomic datasets provide both precision and accuracy in estimating the timescale of placental mammal phylogeny. Proc. Biol. Sci. 279:3491–3500.

dos Reis, M, T. Zhu, and Z. Yang. 2014. The impact of the rate prior on Bayesian estimation of divergence times with multiple Loci. Syst. Biol. 63:555–565.

Drummond, A. J., S. Y. W. Ho, M. J., Phillips, and A. Rambaut. 2006. Relaxed phylogenetics and dating with confidence. PLoS Biol. 4:e88.

Duchêne, S., M. Molak, and S. Y. W. Ho. 2014. ClockstaR: choosing the number of relaxed-clock models in molecular phylogenetic analysis. Bioinformatics 30:1017–1019.

Felice, R. N. and A. Goswami. 2018. Developmental origins of mosaic evolution in the avian cranium. Proc. Natl. Acad. Sci. U. S. A. 115:555–560.

Felsenstein, J. 1973. Maximum-likelihood estimation of evolutionary trees from continuous characters. Am. J. Hum. Genet. 25:471–492.

Felsenstein, J. 1981. Evolutionary Trees from Gene Frequencies and Quantitative Characters: Finding Maximum Likelihood Estimates. Evolution 35:1229–1242.

Felsenstein, J. 1988. Phylogenies and quantitative characters. Ann. Rev. Ecol. Syst. 19:445–471.

Felsenstein, J. 1993. PHYLIP (Phylogeny Inference Package) Version 3.5c. Distributed by the author. Department of Genetics, University of Washington, Seattle.

Felsenstein, J. 2005. Using the quantitative genetic threshold model for inferences between and within species. Philos. Trans. R. Soc. Lond. B. Biol. Sci. 360:1427–1434.

Felsenstein, J. 2012. A comparative method for both discrete and continuous characters using the threshold model. Am. Nat. 179:145–156.

Finarelli, J. A. and A. Goswami. 2009. The evolution of orbit orientation and encephalization in the Carnivora (Mammalia). J. Anat. 214:671–678.

Fisher, R. A. 1919. XV. The correlation between relatives on the supposition of Mendelian inheritance. Trans. R. Soc. Edinburgh 52:399–433.

Foster, C. S. and S. Y. Ho. 2017. Strategies for partitioning clock models in phylogenomic dating: application to the angiosperm evolutionary timescale. Genome Biol. Evol. 9:2752–2763.

Freckleton, R. P. 2012. Fast likelihood calculations for comparative analyses. Methods Ecol. Evol. 3:940–947.

Gavryushkina, A., T. A., Heath, D. T., Ksepka, T. Stadler, D. Welch, and A. J. Drummond. 2017. Bayesian total-evidence dating reveals the recent crown radiation of penguins. Syst. Biol. 66:57–73.

Gavryushkina, A, D. Welch, T. Stadler, and A. J. Drummond. 2014. Bayesian inference of sampled ancestor trees for epidemiology and fossil calibration. PLoS Comput. Biol. 10:e1003919.

Goolsby, E. W. 2016. Likelihood-based parameter estimation for high-dimensional phylogenetic comparative models: overcoming the limitations of “distance-based” methods. Syst. Biol. 65:852–870.

Goswami, A., N. Milne, and S. Wroe. 2011. Biting through constraints: cranial morphology, disparity and convergence across living and fossil carnivorous mammals. Proc. Biol. Sci. 278:1831–1839.

Goswami, A., J. B., Smaers, C. Soligo, and P. D. Polly. 2014. The macroevolutionary consequences of phenotypic integration: from development to deep time. Philos. Trans. R. Soc. Lond. B. Biol. Sci. 369:20130254.

Gower, J. C. 1975. Generalized procrustes analysis. Psychometrika 40:33–51.

Grimm, G. W., P. Kapli, B. Bomfleur, S. McLoughlin, and S. S. Renner. 2015. Using more than the oldest fossils: dating osmundaceae with three Bayesian clock approaches. Syst. Biol. 64:396–405.

Hansen, T. F. 1997. Stabilizing selection and the comparative analysis of adaptation. Evolution 51:1341–1351.

Hasegawa, M., H. Kishino, and T. Yano. 1985. Dating of the human-ape splitting by a molecular clock of mitochondrial DNA. J. Mol. Evol. 22:160–174.

Hasegawa, M, T. Yano, and H. Kishino. 1984. A new molecular clock of mitochondrial DNA and the evolution of hominoids. Proc. Japan Acad. Ser. B. 60:95–98.

Heath, T. A., J. P., Huelsenbeck, and T. Stadler. 2014. The fossilized birth-death process for coherent calibration of divergence-time estimates. Proc. Natl. Acad. Sci. U. S. A. 111:E2957–66.

Ho, S. Y. 2014. The changing face of the molecular evolutionary clock. Trends. Ecol. Evol. 29:496–503.

Ives, A. R., P. E., Midford, and T. Garland. 2007. Within-species variation and measurement error in phylogenetic comparative methods. Syst. Biol. 56:252–270.

King, T., S. Butcher, and L. Zalewski. 2017. Apocrita - High Performance Computing Cluster for Queen Mary University of London.

Lande, R. 1976. Natural selection and random genetic drift in phenotypic evolution. Evolution 30:314–334.

Landis, M. J. and J. G. Schraiber. 2017. Pulsed evolution shaped modern vertebrate body sizes. Proc. Natl. Acad. Sci. U. S. A. 114:13224–13229.

Landis, M. J., J. G., Schraiber, and M. Liang. 2013. Phylogenetic analysis using Lévy processes: finding jumps in the evolution of continuous traits. Syst. Biol. 62:193–204.

Larson-Johnson, K. 2016. Phylogenetic investigation of the complex evolutionary history of dispersal mode and diversification rates across living and fossil Fagales. New Phytol. 209:418–435.

Leaché, A. D., B. L. Banbury, J. Felsenstein, A. N. M. de Oca, and A. Stamatakis. 2015. Short tree, long tree, right tree, wrong tree: new acquisition bias corrections for inferring SNP phylogenies. Syst. Biol. 64:1032–1047.

Lee, M. S. Y. 2016. Multiple morphological clocks and total-evidence tip-dating in mammals. Biol. Lett. 12:20160033.

Lee, M. S. Y., P. M. Oliver, and M. N. Hutchinson. 2009. Phylogenetic uncertainty and molecular clock calibrations: A case study of legless lizards (Pygopodidae, Gekkota). Mol. Phylogenet. Evol. 50:661–666.

Lemey, P., A. Rambaut, J. J. Welch, and M. A. Suchard. 2010. Phylogeography takes a relaxed random walk in continuous space and time. Mol. Biol. Evol. 27:1877–1885.

Lewis, P. O. 2001. A likelihood approach to estimating phylogeny from discrete morphological character data. Syst. Biol. 50:913–925.

Löytynoja, A. and N. Goldman. 2005. An algorithm for progressive multiple alignment of sequences with insertions. Proc. Natl. Acad. Sci. U. S. A. 102:10557–10562.

Löytynoja, A. and N. Goldman. 2008. Phylogeny-aware gap placement prevents errors in sequence alignment and evolutionary analysis. Science 320:1632–1635.

Magallón, S. 2010. Using fossils to break long branches in molecular dating: a comparison of relaxed clocks applied to the origin of angiosperms. Syst. Biol. 59:384–399.

Martín-Serra, A., B. Figueirido, and P. Palmqvist. 2014. A three-dimensional analysis of the morphological evolution and locomotor behaviour of the carnivoran hind limb. BMC Evol. Biol. 14:129.

Matzke, N. J. and A. Wright. 2016. Inferring node dates from tip dates in fossil Canidae: the importance of tree priors. Biol. Lett. 12:20160328.

Meyer, K. and M. Kirkpatrick. 2010. Better estimates of genetic covariance matrices by “bending” using penalized maximum likelihood. Genetics 185:1097–1110.

Nylander, J. A. A., F. Ronquist, J. P., Huelsenbeck, and J. L. Nieves-Aldrey. 2004. Bayesian phylogenetic analysis of combined data. Syst. Biol. 53:47–67.

O’Reilly, J. E., M. dos Reis, and P. C. J. Donoghue. 2015. Dating tips for divergence-time estimation. Trends. Genet. 31:637–650.

Pagel, M. 1994. Detecting correlated evolution on phylogenies: a general method for the comparative analysis of discrete characters. Proc. R. Soc. Lond. B 255:37–45.

Paradis, E., J. Claude, and K. Strimmer. 2004. APE: analyses of phylogenetics and evolution in R language. Bioinformatics 20:289–90.

Parins-Fukuchi, C. 2018a. Bayesian placement of fossils on phylogenies using quantitative morphometric data. Evolution 72:1801–1814.

Parins-Fukuchi, C. 2018b. Use of continuous traits can improve morphological phylogenetics. Syst. Biol. 67:328–339.

Pyron, R. A. 2011. Divergence time estimation using fossils as terminal taxa and the origins of Lissamphibia. Syst. Biol. 60:466–481.

Rannala, B. and Z. Yang. 2007. Inferring speciation times under an episodic molecular clock. Syst. Biol. 56:453–466.

Reeder, T. W., T. M. Townsend, D. G., Mulcahy, B. P., Noonan, P. L. J., Wood, J. W. J. Sites, and J. J. Wiens. 2015. Integrated analyses resolve conflicts over squamate reptile phylogeny and reveal unexpected placements for fossil taxa. PLoS ONE 10:e0118199.

Revell, L. J. and L. J. Harmon. 2008. Testing quantitative genetic hypotheses about the evolutionary rate matrix for continuous characters. Evol. Ecol. Res. 10:311–331.

Ripley, B. D. 1987. Stochastic simulation. Wiley Series in Probability and Statistics John Wiley & Sons, Inc.

Rohlf, F. J. and D. Slice. 1990. Extensions of the Procrustes method for the optimal superimposition of landmarks. Syst. Zool. 39:40–59.

Ronquist, F., S. Klopfstein, L. Vilhelmsen, S. Schulmeister, D. L. Murray, and A. P. Rasnitsyn. 2012. A total-evidence approach to dating with fossils, applied to the early radiation of the Hymenoptera. Syst. Biol. 61:973–999.

Ronquist, F, N. Lartillot, and M. J. Phillips. 2016. Closing the gap between rocks and clocks using total-evidence dating. Phil. Trans. R. Soc. Lond. B. Biol. Sci. 371:20150136.

Schäfer, J. and K. Strimmer. 2005. A shrinkage approach to large-scale covariance matrix estimation and implications for functional genomics. Stat. Appl. Genet. Mol. Biol. 4:Article32.

Schlager, S. 2017. Morpho and Rvcg - Shape Analysis in R: R-Packages for geometric morphometrics, shape analysis and surface manipulations. Pages 217–256 in Statistical shape and deformation analysis (G. Zheng, S. Li, and G. Szekely, eds.). Elsevier.

Schrago, C. G., B. Mello, and A. E. R. Soares. 2013. Combining fossil and molecular data to date the diversification of New World Primates. J. Evol. Biol. 26:2438–2446.

Schraiber, J. G., Y. Mostovoy, T. Y. Hsu, and R. B. Brem. 2013. Inferring evolutionary histories of pathway regulation from transcriptional profiling data. PLoS Comput. Biol. 9:e1003255.

Slater, G. J. 2013. Phylogenetic evidence for a shift in the mode of mammalian body size evolution at the Cretaceous-Palaeogene boundary. Methods Ecol. Evol. 4:734–744.

Slater, G. J., L. J. Harmon, and M. E. Alfaro. 2012. Integrating fossils with molecular phylogenies improves inference of trait evolution. Evolution 66:3931–3944.

Stadler, T. and Z. Yang. 2013. Dating phylogenies with sequentially sampled tips. Syst. Biol. 62:674–688.

Stamatakis, A. 2014. RAxML version 8: a tool for phylogenetic analysis and post-analysis of large phylogenies. Bioinformatics 30:1312–1313.

Tavaré, S., C. R. Marshall, O. Will, C. Soligo, and R. D. Martin. 2002. Using the fossil record to estimate the age of the last common ancestor of extant primates. Nature 416:726–729.

Thorne, J. L., H. Kishino, and I. S. Painter. 1998. Estimating the rate of evolution of the rate of molecular evolution. Mol. Biol. Evol. 15:1647–1657.

Winterton, S. L. and J. L. Ware. 2015. Phylogeny, divergence times and biogeography of window flies (Scenopinidae) and the therevoid clade (Diptera: Asiloidea). Syst. Entomol. 40:491–519.

Wood, H. M., N. J. Matzke, R. G. Gillespie, and C. E. Griswold. 2013. Treating fossils as terminal taxa in divergence time estimation reveals ancient vicariance patterns in the palpimanoid spiders. Syst. Biol. 62:264–284.

Wright, A. M., G. T. Lloyd, and D. M. Hillis. 2016. Modeling character change heterogeneity in phylogenetic analyses of morphology through the use of priors. Syst. Biol. 65:602–611.

Xie, W, P. O. Lewis, Y. Fan, L. Kuo, and M.-H. Chen. 2011. Improving marginal likelihood estimation for Bayesian phylogenetic model selection. Syst. Biol. 60:150–60.

Yang, Z. 1994. Maximum likelihood phylogenetic estimation from DNA sequences with variable rates over sites: approximate methods. J. Mol. Evol. 39:306–14.

Yang, Z. 1995. A space-time process model for the evolution of DNA sequences. Genetics 139:993–1005.

Yang, Z. 2007. PAML 4: Phylogenetic analysis by maximum likelihood. Mol. Biol. Evol. 24:1586–1591.

Yang, Z. and B. Rannala. 2006. Bayesian estimation of species divergence times under a molecular clock using multiple fossil calibrations with soft bounds. Mol. Biol. Evol. 23:212–226.

Zhang, C, T. Stadler, S. Klopfstein, T. A. Heath, and F. Ronquist. 2016. Total-evidence dating under the fossilized birth-death process. Syst. Biol. 65:228–249.

